# Virtual multiplex staining of the pancreatic islets across type 1 diabetes progression using a Schrödinger bridge

**DOI:** 10.64898/2026.04.14.718559

**Authors:** Yu Shen, Won June Cho, Saurabh Joshi, Benjamin Wen, Swarnagouri Naganathanhalli, Maria Beery, Casey Grubel, Arrun Sivasubramanian, Andre Forjaz, Mia P. Grahn, Lucie Dequiedt, Yichen Huang, Kyu Sang Han, Fan Wu, Brian A. Pedro, Laura D. Wood, Tiane Chen, Ralph H. Hruban, Irina Kusmartseva, Mark Atkinson, Denis Wirtz, Ashley L. Kiemen

## Abstract

Classical hematoxylin and eosin (H&E) staining enables review of tissue morphology but lacks information regarding the molecular state of cells. Immunohistochemical (IHC) techniques label specific proteins in tissue, allowing differentiation of relevant structures that may go undetectable in H&E. However, the IHC process is complex, expensive, and time-consuming, especially for multiplex IHC (mIHC) limiting its use in large cohorts. Stain conversion of H&E to IHC using generative artificial intelligence models such as generative adversarial networks (GANs) represent one solution to this problem. However, GANs are unstable during out of distribution sampling and are prone to hallucinations or mode collapse, limiting their accuracy in challenging image conversion tasks. To address this, the field has recently turned to diffusion models. Here, we introduce Schrödinger-bridge for Multiplex ImmunoLabel Estimation (SMILE). Unlike conventional diffusion models that map from source to target through an intermediate Gaussian noise, Schrödinger-bridge diffusion models skip this step and have been shown to better preserve structures during image translation.

To test the performance of SMILE, we generated a large cohort of high-fidelity H&E-mIHC image pairs from pancreatic organ donors, targeting insulin, glucagon, and CD3. Our dataset well-sampled across type-1 diabetes status, pancreas anatomical location, age, and sex. Using this cohort, we demonstrate the superiority of SMILE compared to GANs via a comprehensive evaluation framework incorporating texture, distribution, and antibody-specific metrics, as well as blinded pathologist reviews. We further confirmed the ability of SMILE to generate accurate mIHC images from H&Es generated at an external site, to perform whole slide image conversion, and to generate realistic three-dimensional maps of the pancreatic islets in non-diabetic, auto-antibody positive, and type-1 diabetic donor tissue. Finally, we performed stain conversion of paired H&E to HER2 and Ki67 images in breast cancer, confirming the superiority of SMILE in diverse stain conversion applications. Collectively, this framework provides a scalable pipeline for high-throughput proteomic inference from archival H&Es, providing transformative potential for pancreatic research and digital pathology.

## INTRODUCTION

Hematoxylin and eosin (H&E) stain is the foundation of anatomic pathology, providing sufficient visualization of cellular morphology to distinguish major functional cell types and enabling the majority of routine diagnoses. However, some pathologies require labeling one or multiple proteins via immunohistochemistry (IHC).^1^ For example, in pancreatic pathology, H&E is sufficient to distinguish the exocrine and endocrine compartments, but studying endocrine diseases such as type 1 diabetes (T1D) requires multiplex IHC (mIHC) to distinguish insulin-producing beta cells from glucagon-producing alpha cells and to characterize inflammation around the islets.^2–9^ Despite its diagnostic value, mIHC has practical limitations that constrain its widespread use, requiring time consuming and expensive optimization of each antibody.^10,11^ These constraints are particularly problematic for applications requiring analysis of large tissue cohorts or three-dimensional (3D) tissue reconstruction, where the cost and labor required for mIHC are prohibitive.^12^

To meet this challenge, researchers have demonstrated the potential of virtual stain conversion using deep generative models (**Table S1**). Generative adversarial networks (GANs), particularly pix2pix (p2p),^13^ have dominated stain conversion applications, with numerous studies demonstrating H&E-to-single-plex IHC conversion for various biomarkers.^14–26^ Notably, pyramid-pix2pix (p-p2p)^26^ introduced multi-scale architecture improvements for HER2 labeling in breast cancer. More recently, diffusion models have emerged as a powerful alternative to GANs, achieving state-of-the-art performance in image generation tasks.^27^ Several groups have applied diffusion models to stain conversion tasks, including H&E-to-IHC translation,^28–30^ label-free tissue-to-H&E conversion,^31^ and polarimetric imaging for H&E and fluorescence labeling.^32^ Most applications use conditional diffusion models, which rely on a fixed Gaussian noise prior and perform image conversion using the source image to guide the reverse denoising process. This is conceptually different from GANs, which perform direct image-to-image translation similar to stylistic transfer. Consequently, conditional diffusion models do not always outperform GANs in virtual labeling tasks, as demonstrated by recent work focused on virtual labelling for slide-free microscopy.^33^ Schrödinger bridge models address these limitations by explicitly solving for optimal transport between source and target distributions using stochastic processes rather than mapping from Gaussian noise. The Image-to-Image Schrödinger Bridge (I^2^SB)^34^ uses entropy-regularized optimal transport to preserve structural features during translation, demonstrating superior performance in medical image tasks such as computed tomography super-resolution, denoising and virtual labeling (see details in Supplement 1).^35,36^

Here, we hypothesized that a Schrödinger bridge would achieve superior performance in virtual histology labeling. We present Schrödinger-bridge for Multiplex ImmunoLabel Estimation (SMILE), a workflow utilizing the I^2^SB backbone to perform whole-slide conversion of H&E to mIHC. A compelling application for this technology is in type 1 diabetes (T1D), where accurate detection of hormone-producing cells in the pancreatic islets of Langerhans is critical for studying the heterogeneous patterning of disease progression. Type 1 diabetes (T1D) is an autoimmune condition characterized by immune mediated destruction of the insulin-producing beta cells, leading to impaired blood glucose regulation. With increasing incidence worldwide,^37^ T1D represents not only one of the most common chronic childhood diseases, but also one that is increasingly diagnosed in adults.^38^ Many individuals harbor auto-antibodies prior to overt T1D onset, yet the spatial patterning of insulin-producing beta cell loss and CD3+ T cell infiltration remains poorly understood.^39^ One major factor limiting study of T1D is the scarcity of high-quality tissue specimens, as T1D individuals rarely undergo pancreatic resection. This void has recently been filled by an emerging number of programs, such as the Network for Pancreatic Organ donors with Diabetes (nPOD) and Human Islet Research Network (HIRN), seeking to meet this need through collection of high-quality tissues obtained from organ donors.^40,41^

We used SMILE to virtually convert H&E images of human pancreas tissue to mIHC with discrete labels for insulin, glucagon, and CD3 (**Fig 1**). As part of this work, we generated a uniquely diverse and high-quality cohort of paired image tiles. This cohort contains organ donor pancreas tissue well sampled across age, sex, T1D status, and location within the pancreas. With this dataset, we demonstrate the superior performance of SMILE compared to p2p and p-p2p in stain conversion of H&E to mIHC, assessed using standard metrics in the field and blinded pathologist review. We evaluated SMILE’s performance on a testing set generated at an external site. Finally, we demonstrate the utility of this model through generation of 3D, cm^3^-scale maps of pancreas tissue, revealing novel insights into spatial patterning with T1D onset. In summary, SMILE introduces a high-fidelity paired H&E and mIHC dataset for benchmarking label conversion models and supports the superiority of Schrödinger bridge models compared to GANs in histology applications.

**Figure 1:**
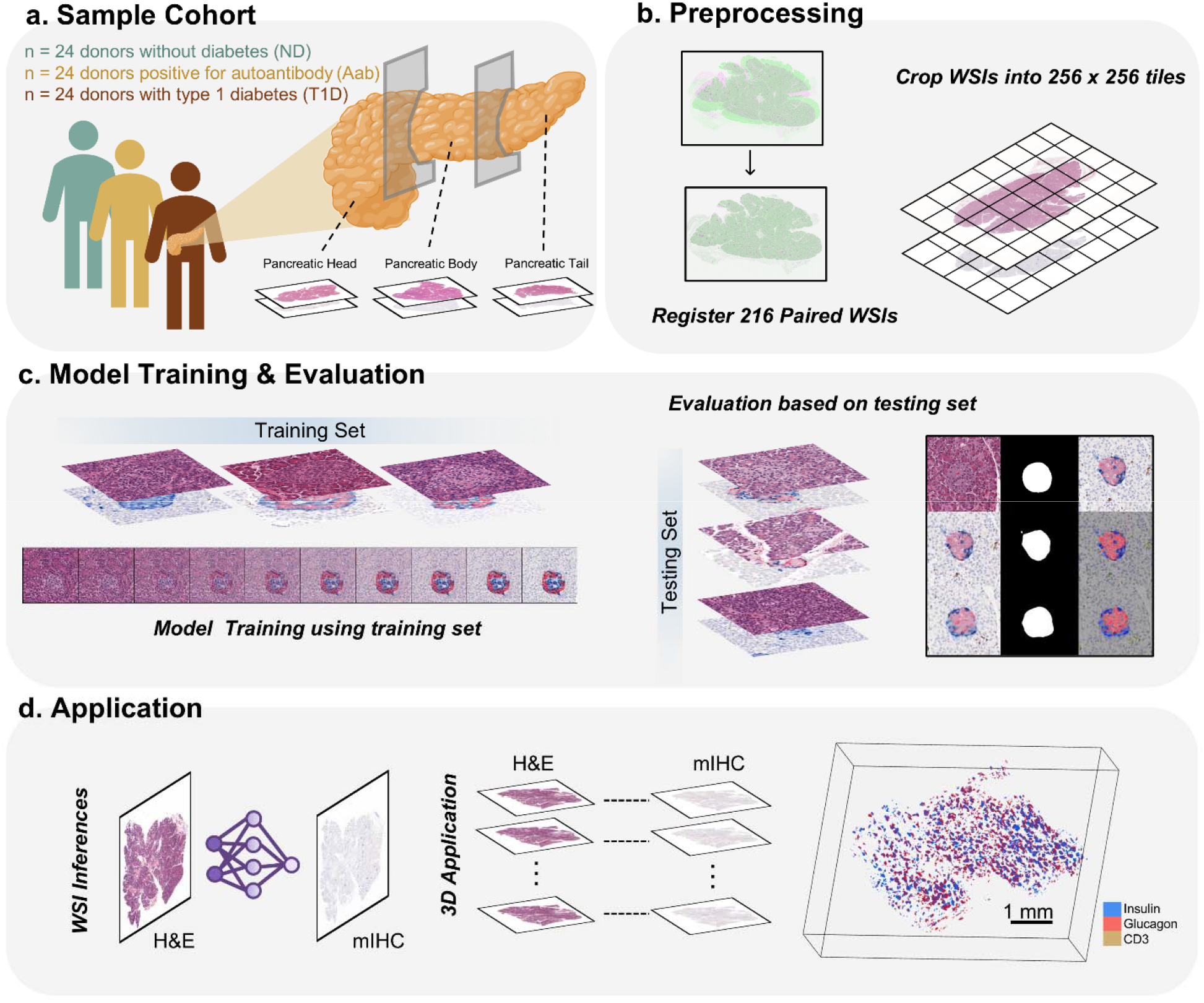
Virtual, multiplex labeling of pancreatic islets of Langerhans. **(a)** Collection of 216 paired H&E and mIHC labeled sections from 72 pancreatic donors from 3 pancreatic locations (pancreas head, body, and tail) with 3 health conditions: donors without diabetes (ND), donors positive for autoantibody (Aab+), and donors with type 1 diabetic (T1D). mIHC image labels for insulin (red), glucagon (blue), and CD3 (brown). **(b)** A computational workflow for nonlinear, 20x-resolution image registration and tiling for model training. **(c)** Virtual labeling models trained on paired image tiles and tested for their ability to generate islets of Langerhans with realistic morphological and molecular features, as tested quantitatively and via blinded pathologist review. **(d)** Application of virtual labeling workflow to generate three-dimensional maps of pancreatic islets.

## MATERIALS AND METHODS

### Pancreatic histology sample collection and digitization

This retrospective study was approved by the Johns Hopkins University and the University of Florida Institutional Review Boards. Samples used for training were collected from organ donors consented for research as part of the Network for Pancreatic Organ donors with Diabetes (nPOD). For each resected pancreas, tissue was collected from the pancreatic head, body, and tail, formalin-fixed and paraffin embedded. Two 4µm thick sections were cut from each sample, for a total of six sections per donor. One slide of each pair was stained with H&E. The other section was labeled with antibodies for insulin (in red), glucagon (in blue), and CD3+ T cells (in brown) and counterstained with hematoxylin. Sections were digitized at 20x resolution using either an Aperio CS2 or Aperio GT450 slide scanner (Leica Biosystems). Samples used for 3D reconstruction were similarly collected by nPOD and consisted of tissue from one donor without diabetes (ND), one donor positive for GADA, ZnT8A, and IA2A (Aab+), and type 1 diabetic (T1D) donor, sampled from the pancreas body. For each sample, 100 4µm thick H&E-stained serial sections were collected, with a spacing of 12µm between consecutive slides. Sections from ND and T1D blocks were digitized at 20x magnification using Aperio CS2 slide scanner (Leica Biosystems), while the sections from Aab+ were digitized at 40x magnification using Aperio GT450 slide scanner (Leica Biosystems). Four external validation samples were collected at Johns Hopkins University (JHU). Pancreatic tissue from two donors without diabetes and two donors with T1D who underwent pancreatic resection following diagnosis of pancreatic ductal adenocarcinoma or severe pancreatitis were retrospectively collected. Grossly normal pancreas tissue was collected from the surgical margin, formalin-fixed, and paraffin embedded. Two 4 - µm thick histological sections were collected from each sample and stained using the same H&E and triple-IHC stain as was used for the initial cohort. Sections were digitized at 20x resolution using a CS2 slide scanner (Leica Biosystems).

### Paired H&E to mIHC image registration

The Openslide algorithm was used to save reduced resolution copies of the 20x images (∼0.5 µm per pixel) at pseudo-10x (1 µm per pixel) and pseudo-2x (5 µm per pixel) resolution.^42^ Logical masks of tissue-containing pixels were created for the 2x H&E and mIHC images through green-channel intensity thresholding and eliminating pixels with low red-green-blue (RGB) standard deviation. Following this, CODA nonlinear image registration was applied to each image pair within a common coordinate system.^2^ The accuracy of image alignment for each pair was validated through manual inspection using Fiji ImageJ. Subsequently, the nonlinear displacement fields computed from the 2x TIF images were applied to the 10x TIF images, resulting in the creation of high-resolution aligned image pairs. Additionally, a script was generated to create high-resolution registered 20x image pairs in OME-TIF format, enabling users to view and save tiles from the cohort at native resolution.

### Generation and quality control of registered training tile image pairs

We developed an algorithm for the rapid generation of high-quality training tiles, balancing tiles containing and not-containing islets of Langerhans. The registered 2x image pairs were imported into MATLAB 2024b, where they were overlaid to facilitate enhanced visualization of the pancreatic anatomy in both H&E and mIHC. The user was prompted to manually select islets in each image. Following selection, the algorithm automatically cropped and exported high-resolution 10x tiles of dimensions 256 × 256 × 3 pixels in PNG format. To ensure spatial accuracy, we implemented a quality control step to manually inspect each tile pair for misalignment and anatomical consistency. Additionally, the algorithm sampled non-islet containing regions to ensure training on histological regions of vasculature, fat, pancreatic acini, pancreatic ducts, and background. The final set of quality-controlled tile pairs were subsequently utilized for model training and evaluation.

### PIX2PIX model training

The pix2pix (p2p) model was implemented using the MATLAB Deep Learning Toolbox (https://github.com/matlab-deep-learning/pix2pix).^13^ The generator employed a U-Net architecture with skip connections operating on 256 × 256-pixel inputs, while the discriminator used a 70 × 70 PatchGAN architecture that classifies overlapping image patches as real or virtual. The model was trained using the default hyperparameters: an Adam optimizer with a learning rate of 0.0002 and momentum parameter β_1_ = 0.5, a batch size of 1, and the combined conditional GAN and L_1_ reconstruction loss with λ = 100. Training proceeded for 200 epochs, with the learning rate held constant for the first 100 epochs and linearly decayed to zero over the remaining 100 epochs. The 39,160 quality-controlled H&E-IHC tile pairs (256 × 256 × 3 pixels) were used for training, with H&E tiles as inputs and the corresponding registered mIHC tiles as targets. Mathematical details of the cGAN objective and L_1_ loss are provided in Supplement 1. At inference, a single forward pass through the trained generator produces the virtual mIHC tile from a given H&E input; the discriminator is not used during generation.

### Pyramid-PIX2PIX model training

The pyramid-pix2pix (p-p2p) model was implemented using the publicly available PyTorch codebase (https://github.com/bupt-ai-cz/BCI).^26^ The generator used a ResNet architecture with nine residual blocks, while the discriminator employed the same 70 × 70 PatchGAN as in p2p. The p-p2p model extends the standard pix2pix framework with a multi-scale pyramid loss: the adversarial and L_1_ losses are computed at full resolution, after which the ground-truth and generated images are progressively smoothed with a Gaussian kernel (3 × 3, σ = 1) and downsampled, and additional L_1_ losses are computed at each pyramid level to account for spatial misalignments between adjacent tissue sections. The model was trained using default hyperparameters: an Adam optimizer with a learning rate of 0.0002 and β_1_ = 0.5, a least-squares GAN (LSGAN) loss, a batch size of 2, and batch normalization throughout. Training proceeded for 200 epochs, with the learning rate held constant for the first 100 epochs and linearly decayed to zero over the remaining 100 epochs. The same 39,160 tile pairs were used for training. Further mathematical details of the GAN framework are provided in Supplement 1. At inference, a single forward pass through the trained generator produces the virtual mIHC output; the pyramid loss computation and discriminator are not used during generation.

### SMILE model training using I^2^SB backbone

The SMILE workflow includes training the Image-to-Image Schrödinger Bridge (I^2^SB) model, which was implemented using the official PyTorch codebase (https://github.com/NVlabs/I2SB).^34^ The network architecture employed the Ablated Diffusion Model (ADM) U-Net, a residual U-Net with attention layers originally designed for diffusion-based image generation. The diffusion process was discretized into 1,000 timesteps with a noise schedule parameterized by β_max_ = 0.3. Training was performed using the Adam optimizer with a learning rate of 5 × 10^−5^, decayed by a factor of 0.99 per epoch. An exponential moving average (EMA) with a decay rate of 0.99 was applied to the model weights during training for stable inference. The model was trained with a global batch size of 128 and a microbatch size of 1 for 100 epochs on the same 39,160 tile pairs (256 × 256 × 3 pixels). All other hyperparameters followed the defaults of the I^2^SB codebase. The theoretical framework of the Schrödinger bridge formulation and the SMILE training objective are detailed in Supplement 1. At inference, SMILE generates virtual mIHC tiles through an iterative sampling process that begins directly from the source H&E tile rather than from Gaussian noise, as in standard diffusion models. Starting from the input H&E tile, the trained network iteratively refines the image over a series of denoising steps toward the target mIHC distribution, producing the final virtual mIHC tile. The number of function evaluations (NFE) controls the number of intermediate denoising steps during this reverse process, governing a trade-off between computational cost and output quality. Unlike the single-pass deterministic generation of GAN-based models, SMI sampling is stochastic and iterative.

### Whole-slide-image (WSI) immunolabel conversion

To perform whole-slide inference, each gigapixel H&E image was first split into smaller, uniformly sized 256 × 256 tiles, with inclusion restricted to tiles demonstrating sufficient tissue coverage as determined by a binary tissue mask. Each valid tile was preprocessed and passed through the deep learning model in batches, ensuring efficient memory usage. The model’s output tiles were subsequently stitched back into the original WSI layout using a weighted average in overlapping regions to correct for borders and edge effects. This tile-based approach enabled high-resolution, end-to-end inference over entire WSIs, preserving spatial context and allowing for efficient computation on very large pathological images where direct inference is not feasible due to hardware constraints.

### Optimize WSI inference speed by reducing image precision

The original I^2^SB architecture was built and tested with 16-bit images. To improve inference speed, SMILE was tested not only at the original 32-bit precision but also at 16-bit precision (**Fig S1a**). We utilized exponential moving average (EMA) with cuDNN version >2.2 to enable fp 16 network weights for faster inference. We inferred the same WSI using SMILE at fp 16, SMILE at fp 32, and p2p at fp 32 (**Fig S1b**). We quantified the inference time as well as performed image quality analysis, demonstrating that that reducing image precision can significantly reduce inference time while maintaining image quality.

### Label mask generations for quantitative evaluations

We developed an algorithm to determine the optimal hue-saturation-value (HSV) range for each of the four colors of interest within four-plex mIHC images. The algorithm randomly selected nine image tiles (**Fig S2a**), each containing four relevant labels: hematoxylin (navy blue), insulin (red), glucagon (bright blue), and CD3 (brown), and concatenated them into a 3 × 3 mosaic image. Utilizing the OpenCV and matplotlib libraries in Python, we represented each pixel of the mosaic in 3D HSV color space (**Fig S2b**), allowing users to identify the HSV ranges for each label through a browser-accessible HTML file. Using the saved HSV range, we segmented the mIHC images into one of five categories: hematoxylin-positive, insulin-positive, glucagon-positive, CD3-positive, or background. To validate this algorithm, the user manually annotated 1576-pixels of insulin regions, 1561-pixels of glucagon regions, and 1481-CD3 regions using Aperio ImageScope on the mosaic image. Comparing the manual annotations (ground truth) and the HSV algorithm predictions on these regions, we quantified the algorithm accuracy (**Fig S2c**).

### Islet mask generations for quantitative evaluations

Islet masks were generated from H&E and mIHC tiles by adapting the CODA segmentation model.^7^ We selected 78 H&E tiles and 52 mIHC tiles and performed annotations of “islet” and “non-islet” regions using Aperio ImageScope. From these annotations we trained two models: one for identifying islet in H&E and another for identifying islet in mIHC and compared their performance ground truth annotations on independent images (**Fig S3**). The resulting islet mask is a labeled image containing numerical indices 1 and 2, corresponding to islet and non-islet regions, respectively. These masks were used for evaluating islet morphology between ground-truth and virtual tiles.

### Quantitative texture and distribution-based validation metrics

After sampling the virtual mIHC-labeled images from the test H&E dataset using the three trained virtual labeling models, seven different texture- and distribution-based metrics were used to evaluate performance. As shown in Table 1, we selected some of the most commonly used quantitative metrics from previous works (see metrics used in previous works listed in Table S1), in addition to our own domain-specific metrics designed to capture islet morphological and molecular features. The first three metrics are texture-based metrics that evaluate pixel-level similarity, while the latter four are distribution-based metrics that assess feature-level distributional similarity.

#### Texture-based metrics

(1) Structural similarity index measure (SSIM):^43^ A metric that combines the luminance, contrast, and structure of the image pair, calculated based on pixel mean, variance, and covariance along the x and y axes. SSIM is widely used to evaluate the perceptual fidelity of image pairs. (2) Complex-wavelet structural similarity index measure (CW-SSIM):^44^ An extension of SSIM calculated in the complex wavelet domain, providing robustness to small geometric distortions such as translation and rotation that may occur between adjacent tissue sections. (3) Peak signal-to-noise ratio (PSNR): A classical metric that calculates the ratio between the maximum pixel value and the mean-squared error between the image pair, commonly used to evaluate image reconstruction and compression quality.

#### Distribution-based metrics

(4) Fréchet inception distance (FID):^45^ A metric that uses a pretrained Inception network to extract feature embeddings from image sets, models them as multivariate Gaussian distributions, and computes the 2-Wasserstein distance between distributions. While FID is widely used to evaluate the fidelity and diversity of generated images, the metric requires large sample sizes for reliable estimation. (5) CLIP-Maximum mean discrepancy (CMMD):^46^ A metric proposed to address FID’s limitations in representation quality and sample efficiency by using pretrained CLIP embeddings and computing the maximum mean discrepancy between feature distributions. CMMD provides more robust evaluation with smaller sample sizes; we additionally report FID for comparison with previous works. (6) Virchow 2 Maximum Mean Discrepancy (VMMD): Analogous to CMMD, this metric replaces the general-purpose CLIP model with Virchow 2,^47^ a state-of-the-art pathology vision foundation model trained on millions of histopathology whole slide images. By computing the maximum mean discrepancy between Virchow 2 embeddings, VMMD provides a domain-specific evaluation of distributional similarity that is better suited for histopathology image assessment. (7) UNI2 embedding similarity: Using UNI2,^48^ another pathology foundation model, we compute the cosine similarity between the virtually labels and ground-truth mIHC image embeddings. This metric directly quantifies the histological similarity between virtual and real images in a pathology-specific feature space.

### Quantitative validation metrics specific to pancreatic endocrine pathology

Insulin, glucagon, and CD3 labels are the primary components visible in our mIHC images, distinguished by their respective colors. To qualitatively assess model performance and correlations between ground-truth mIHC and virtual mIHC tiles, we compared the morphological shape of the ground truth H&E to the virtual mIHC and compared the islet label composition between the ground truth mIHC and the virtual mIHC. The islet masks generated by the CODA segmentation model were used to compute intersection over union (IOU) between ground-truth and virtual mIHC. A higher IOU indicates greater morphological similarity. The compositional accuracy of virtual mIHC images compared to ground-truth mIHC images is determined using extracted label masks derived from HSV color space. By utilizing label masks for each label component, we evaluated both regional and global differences in each label between ground-truth and virtual images.

### Qualitative validation via blinded pathologist review

To compare each model’s ability to generate convincing virtual mIHC-labeled images, we developed a graphic user interface (GUI) to randomly load tiles generated from three deep learning models and asked two pathologists to rank the model performance based on the histological quality and consistency the ground-truth image (**Fig S4**). Utilizing the PyQt5 library in Python, we established the GUI to randomly select 50 virtual tiles generated by the SMILE, p2p, and p-p2p models. In the first assessment, pathologists evaluated the pathological quality by ranking the virtual tiles from three models in orders of 1, 2, and 3, where a rank of 1 corresponds to the best quality and receives 3 points, a rank of 2 corresponds to the second-best quality and receives 2 points, and a rank of 3 corresponds to the lowest quality and receives 1 point. In the second task, the corresponding ground-truth H&E and mIHC tiles were presented, and the pathologist ranked the virtual tiles based on their level of consistency with respect to the ground-truth images, using the same ranking system as in the first assessment. Ultimately, we averaged the scoring results from both pathologies for a total of 100 randomly selected tiles.

### Three-dimensional (3D) tissue and label reconstructions of virtual mIHC sections

We performed WSI label conversion of serial H&E images from one ND donor, one Aab+ donor, and one T1D donor unseen during training to generate serial virtual mIHC image stacks. The Openslide algorithm described above was used to generate pseudo-2x and pseudo-10x images from the high-resolution scans. The CODA nonlinear registration algorithm described above was used to calculate registration on the serial images, and the HSV segmentation method described above was used to discriminate insulin+, glucagon+, and CD3+ regions from the virtual images. Volumetric matrices were generated at a resolution of 4 µm × 4 µm × 12 µm and rendered using the image processing toolbox in MATLAB 2024b.

CODA semantic segmentation algorithm was applied on H&E slides. Eight distinct tissue classes were manually annotated across 18 H&E slides representing three donor samples using Aperio ImageScope software. The model training was accomplished using MAATLAB 2024b, achieving a segmentation accuracy of 94.9 %. All H&E slides were subsequently segmented into tissue-labeled images with a resolution of 1 µm per pixel. 3D reconstruction was achieved through image registration of segmented tissue masks for each tissue blocks, generating a volumetric matrix with a resolution of 4 µm × 4 µm × 12 µm and rendered using the image processing toolbox.

Following previously published standards defining endocrine cell clusters and pancreatic islets,^49^ we discarded insulin and glucagon positive objects smaller than 3,000 µm^3^. Objects between 3,000 - 10,000 µm^3^ were termed endocrine cell clusters, and objects larger than 10,000 µm^3^ were termed pancreatic islets. Additionally, only signals co-localizing with the H&E-derived islet mIHC label mask were retained for quantification. For each endocrine cluster and pancreatic islet, we calculated the insulin composition, glucagon composition, and composition of CD3 in the 50 µm^3^ surrounding each object.

## RESULTS

### A Novel Dataset for H&E to mIHC Label Conversion to Image T1D Progression

Pancreatic tissue was retrospectively collected from 72 organ donors diverse in sex, age, race, and T1D status (**Table S1**). As all of these demographic features are known to influence pancreatic islet size, distribution, and composition,^50–54^ this cohort design ensures collection of images with diverse presentation of islets. Tissue sections each were collected from the pancreatic head, body, and tail of each donor and stained using H&E and mIHC for insulin-glucagon-CD3. The mIHC images enabled easy recognition of the pancreatic islets of Langerhans in the ND, Aab+, and T1D pancreas, even where the islets could at times be difficult to distinguish in H&E. Seen through visual inspection, the samples without diabetes contained greater composition of insulin-producing beta cells (in red) than the Aab+ and T1D samples, while the Aab+ and T1D samples contained greater composition of glucagon-producing alpha cells (in blue) and CD3+ T cells (in brown) than the ND samples, conforming to known beta cell loss and insulitis with onset of T1D.^55^

We applied CODA nonlinear image registration to H&E and mIHC WSI pairs. Following registration of low-resolution images, we developed a workflow to apply the registration transforms to the native-resolution WSIs, generating registered, native resolution OME-TIF files. We visually inspected these results and removed four WSI pairs due to extensive tissue folding or large anatomical mismatch between the H&E and mIHC, leaving a total of 212 WSI pairs for model construction. We randomly selected 6 WSI pairs for model testing, consisting of 2 pairs each from ND, Aab+, and T1D donors, leaving 206 WSI pairs for model training.

We noted that islets of Langerhans and CD3+ T cells were minority populations in the pancreatic tissue. This observation was confirmed by previous works that estimate that islets comprise 1-5% of pancreas mass.^56,57^ Therefore, random sampling of the image pairs would result in training tiles that were dramatically imbalanced between negatively and positively labeled structures, leading to poor model performance. To balance the ratio of positive and negative signals, we designed an algorithm to rapidly generate high-quality tiles centered around the islets of Langerhans as well as other pancreatic tissue structures. We manually reviewed the overlay of each cropped tile pair and removed pairs deemed low quality due to imperfect registration or large change in pancreatic anatomy across serial sections. Our quality-controlled training dataset consisted of 39,160 tile pairs. The testing datasets were similarly constructed, for a final size of 1,293 tile pairs in the internal testing dataset. Our final cohort and tile metrics are presented in **Fig 2**, with per WSI pair and per tile demographic distributions provided.

**Figure 2:**
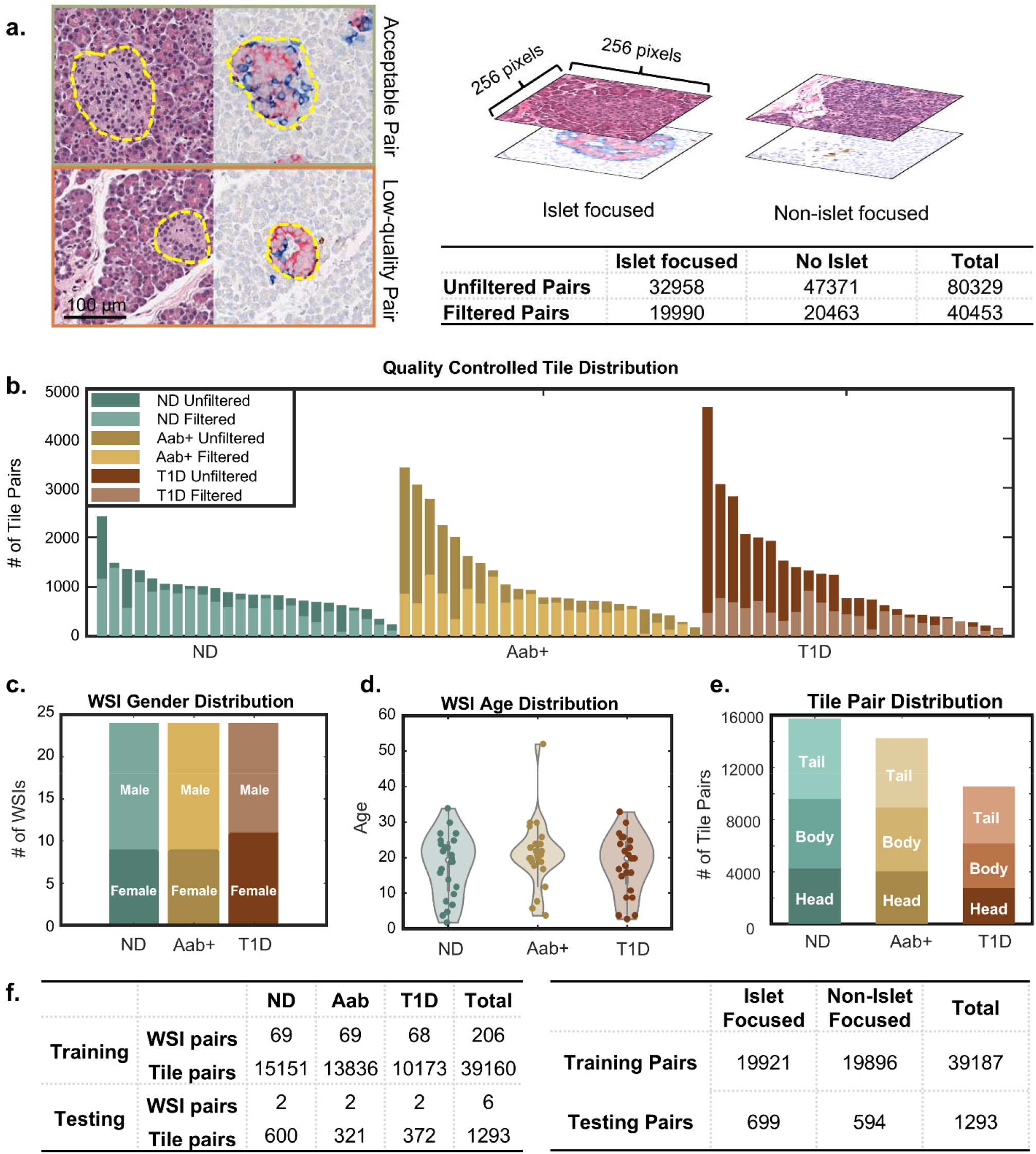
A curated cohort for histology image conversion. **(a)** Tiles were generated at a size of 256 × 256 pixels, with roughly 50% chosen to contain an islet and 50% chosen not to contain an islet. Tiles were manually reviewed to remove poorly aligned pairs. Sample high-quality and low-quality image pairs shown, with the dashed outline highlighting the islets. The table displays total numbers of raw and filtered tiles. **(b)** Distribution of all tiles before and after filtering. **(c - e)** Bar and violin plots presenting donor distribution by diabetes status, age, and sex. **(f)** Summary statistics describing the final training and testing distribution by diabetes status and the number of islet-focused and non-islet-focused tiles.

### SMILE model training and WSI inference

Using the 39,160 quality-controlled tile pairs, we trained three generative models for H&E-to-mIHC conversion: pix2pix (p2p), pyramid-pix2pix (p-p2p), and SMILE. The p2p and p-p2p models represent the current standard for paired virtual labeling, employing GAN-based architectures that perform direct image-to-image translation through adversarial training. The SMILE workflow uses an alternative approach based on the Schrödinger bridge framework, which learns to optimally transport the H&E distribution to the mIHC distribution through a stochastic diffusion process that begins directly from the source image rather than Gaussian noise (see Supplement 1 for technical details). All three models were trained using their respective default hyperparameters as described in the Materials and Methods. Following training, each model was used to convert the 1,293 held-out testing H&E tiles into virtual mIHC tiles. Visual inspection of the virtual outputs revealed that all three models successfully learned to generate the mIHC color palette, insulin (red), glucagon (blue), and CD3 (brown), from the H&E inputs, though notable differences in image quality and labeling fidelity were apparent across models (**Fig 3a**). For evaluation purposes, we used HSV color intensity to segment insulin+, glucagon+, CD3+, and antibody-pixels from the ground truth and virtual mIHC images and trained semantic segmentation models to segment the islet morphology in all ground truth and virtual mIHC images (**Fig 3b**). These masks allowed us to compare the islet shape and label positivity between ground truth and virtual images.

**Figure 3:**
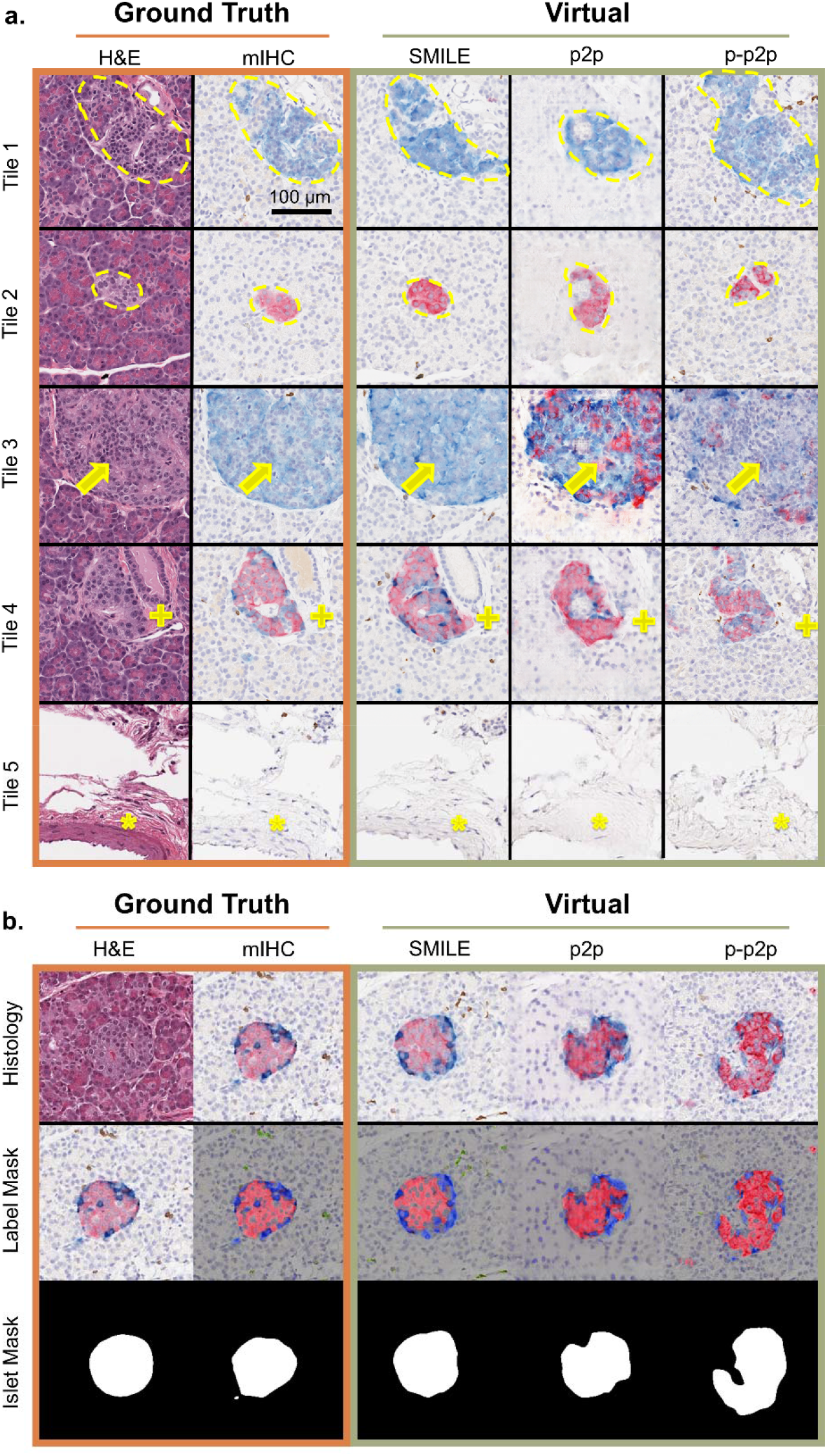
Visual comparison of ground truth and virtual tiles. **(a)** Mosaic presenting ground truth histology (GT H&E, GT mIHC) and virtual mIHC generated using three models (SMILE, p2p, p-p2p) of five representative testing tiles. Dashed line outlines islet morphology in Tile 1 and Tile 2. Tile 3: yellow arrow pointing to islet labeling, where the SMILE image most closely matches the GT mIHC. Tile 4: plus sign indicating a pancreatic duct adjacent to an islet. Only the SMILE model correctly generates the duct. Tile 5: Asterisk indicates an artery. Only the SMILE model correctly generates the artery. **(b)** Representative images of GT H&E, GT mIHC, virtual mIHC, label masks (to segment the mIHC signal), and islet masks (to segment the islet morphology), used to validate the virtually labeled images.

For whole-slide inference, each trained model was applied in a tile-based fashion: the WSI was partitioned into 256 × 256-pixel tiles, each tile was independently converted to virtual mIHC, and the resulting tiles were stitched back into the full WSI coordinate space using weighted averaging to mitigate border artifacts (see Materials and Methods). To systematically evaluate and optimize model performance before final benchmarking, we first conducted a hyperparameter search specific to SMILE, as described in the following section.

### Identification of optimal SMILE model weight and number of functional equations (NFE) used for inference

Training and testing large diffusion models are computationally heavy. In I^2^SB, the number of functional equations (NFE) can be reduced with negligible tradeoff of performance because of the inherent optimal transport framework, the number of inferencing timesteps can be low and the optimal path can still be approximated well. We obtained ten model weights at epochs 10, 20, 30, 40, 50, 60, 70, 80, 90, and 100. At each model weight, we inferred 1,293 testing H&E tiles into virtual mIHC tiles using NFE values of 16, 32, and 62. In total, we generated model outputs for 30 combinations of model weights and NFEs (**Fig S5a**). To identify the optimal training weight and NFE for virtual labeling, we employed quantitative evaluations methods based on label masks and islet masks. We computed and plotted intersection over union (IOU), regional label difference (RSD), global insulin label difference (GID), and global glucagon label difference (GGD) for each combination (**Fig S5b-4e**). We then calculated a combined score for each combination using min-max normalization. Detailed calculations are documented in Supplementary 2. Ultimately, we identified the optimal weight for inference to be the one obtained at 80 epochs with NFE set to 32 (**Fig S5f**), and the inference time for each tile is approximately 3.5 seconds per tile (**Fig S5g**).

### SMILE outperforms current state-of-the-art models in virtual labeling of the pancreatic islets as assessed through various performance metrics

We compared the performance of SMILE to p2p and p-p2p based using thirteen performance metrics chosen based on their ability to evaluate similarity between adjacent sections that may be structurally dissimilar (**Table 3**). These metrics are divided into texture-based metrics: structural similarity index measure (SSIM), complex-wavelet structural similarity index measure (CW-SSIM), and peak signal-to-noise ratio (PSNR); four distribution-based metrics: Fréchet inception distance (FID), CLIP-maximum mean discrepancy (CMMD), Virchow 2 maximum mean discrepancy (VMMD), and UNI2 embedding similarity; four label-based metrics: intersection over union (IOU), regional label differences (RSD), global insulin differences (GID), and global glucagon differences (GGD); and two pathologist review questions (PATH Q1 and Q2). Information about these metrics can be found in the Materials and Methods section.

**Table 3:**
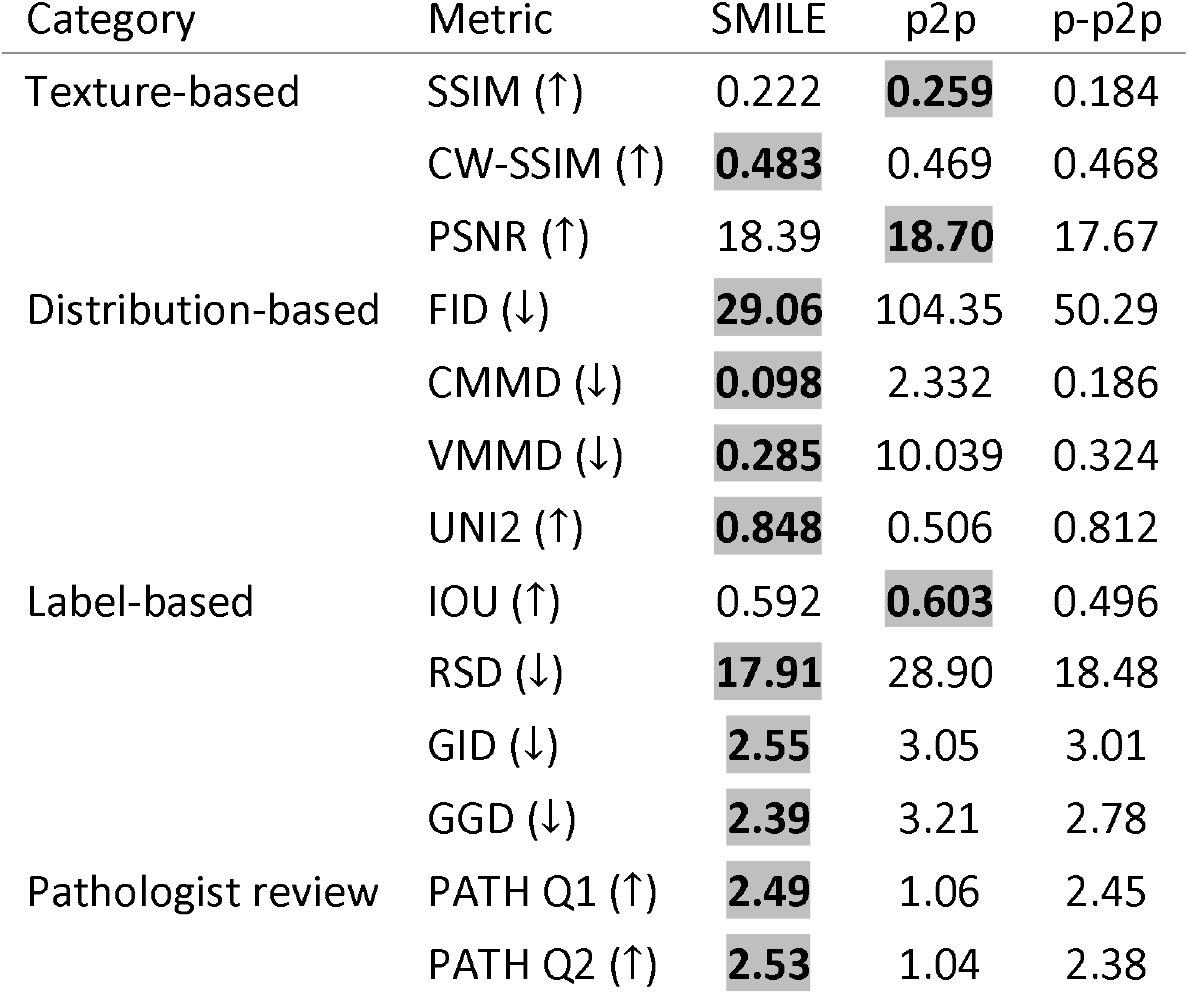
Quantitative evaluation of virtual labeling models. Arrows indicate whether higher (↑) or lower (↓) values are better. Bold values indicate the best performance for each metric. Texture-based metrics evaluate pixel-level similarity, distribution-based metrics assess feature-level distributional similarity using pretrained neural networks, and label-based metrics quantify morphological and compositional accuracy of the virtual mIHC labels.

| Category | Metric | SMILE | p2p | p-p2p |
| --- | --- | --- | --- | --- |
| Texture-based | SSIM (↑) | 0.222 | **0.259** | 0.184 |
|  | CW-SSIM (↑) | **0.483** | 0.469 | 0.468 |
|  | PSNR (↑) | 18.39 | **18.70** | 17.67 |
| Distribution-based | FID (↓) | **29.06** | 104.35 | 50.29 |
|  | CMMD (↓) | **0.098** | 2.332 | 0.186 |
|  | VMMD (↓) | **0.285** | 10.039 | 0.324 |
|  | UNI2 (↑) | **0.848** | 0.506 | 0.812 |
| Label-based | IOU (↑) | 0.592 | **0.603** | 0.496 |
|  | RSD (↓) | **17.91** | 28.90 | 18.48 |
|  | GID (↓) | **2.55** | 3.05 | 3.01 |
|  | GGD (↓) | **2.39** | 3.21 | 2.78 |
| Pathologist review | PATH Q1 (↑) | **2.49** | 1.06 | 2.45 |
|  | PATH Q2 (↑) | **2.53** | 1.04 | 2.38 |

For texture-based metrics, p2p achieved the highest SSIM and PSNR, while SMILE performed best on CW-SSIM. Of note, the differences among models were relatively modest. In contrast, SMILE demonstrated substantial superiority across all four distribution-based metrics. For FID, SMILE achieved a score of 29.06 compared to 50.29 for p-p2p and 104.35 for p2p, representing a 3.6-fold improvement. The gap was even more pronounced for CMMD, where SMILE outperformed p-p2p by 1.9-fold and p2p by 23.8-fold. Similarly, VMMD showed SMILE substantially outperforming p-p2p and p2p. For UNI2 embedding similarity, SMILE achieved the highest score, followed by p-p2p and p2p. SMILE achieved the best performance in three of four label-based metrics: RSD, GID, and GGD. However, p2p achieved a marginally higher IOU compared to SMILE and p-p2p. From the pathologist reviews, SMILE demonstrated superior performance in both assessment tasks. The results of PATH Q1 indicate that the virtual tiles generated using the SMILE model best replicate traditional mIHC histological features. Similarly, PATH Q2 reveals that the virtual tiles produced by SMILE model are the most consistent with the GT images. This suggests that the virtual mIHC closely resembles the expected appearance of GT mIHC labeling based on human perception.

Overall, SMILE outperforms the GAN-based models on the majority of evaluation metrics, particularly excelling in distribution-based metrics that capture global perceptual quality. The p2p model excelled for quantitative metrics where discrete pixels are compared but faltered where spatial continuity or realism is required.

### 3D spatial reconstruction of insulin and glucagon

As a use case to determine the ability of the SMILE to facilitate new insights into T1D spatial biology, we inferred mIHC images in 3D from serial H&E-stained pancreas images (**Fig 4a**). We generated roughly 100 serial H&E images each from ND, Aab+, and T1D organ donors using a new cohort unseen during model training or testing. Using the CODA algorithm, we registered the serial H&E images into a semi-continuous stack, then applied the resulting transformation matrices to the inferred mIHC images. Using our HSV segmentation algorithm we determined label positivity on each of the virtual mIHC images and combined these to generate labelled 3D maps. Within these reconstructions we could determine tissue volume and islet number (**Fig 4b**). Using previously described thresholds to categorize the endocrine cells of the pancreas,^49^ we discarded insulin and glucagon positive objects smaller than 3,000 µm^3^.

**Figure 4:**
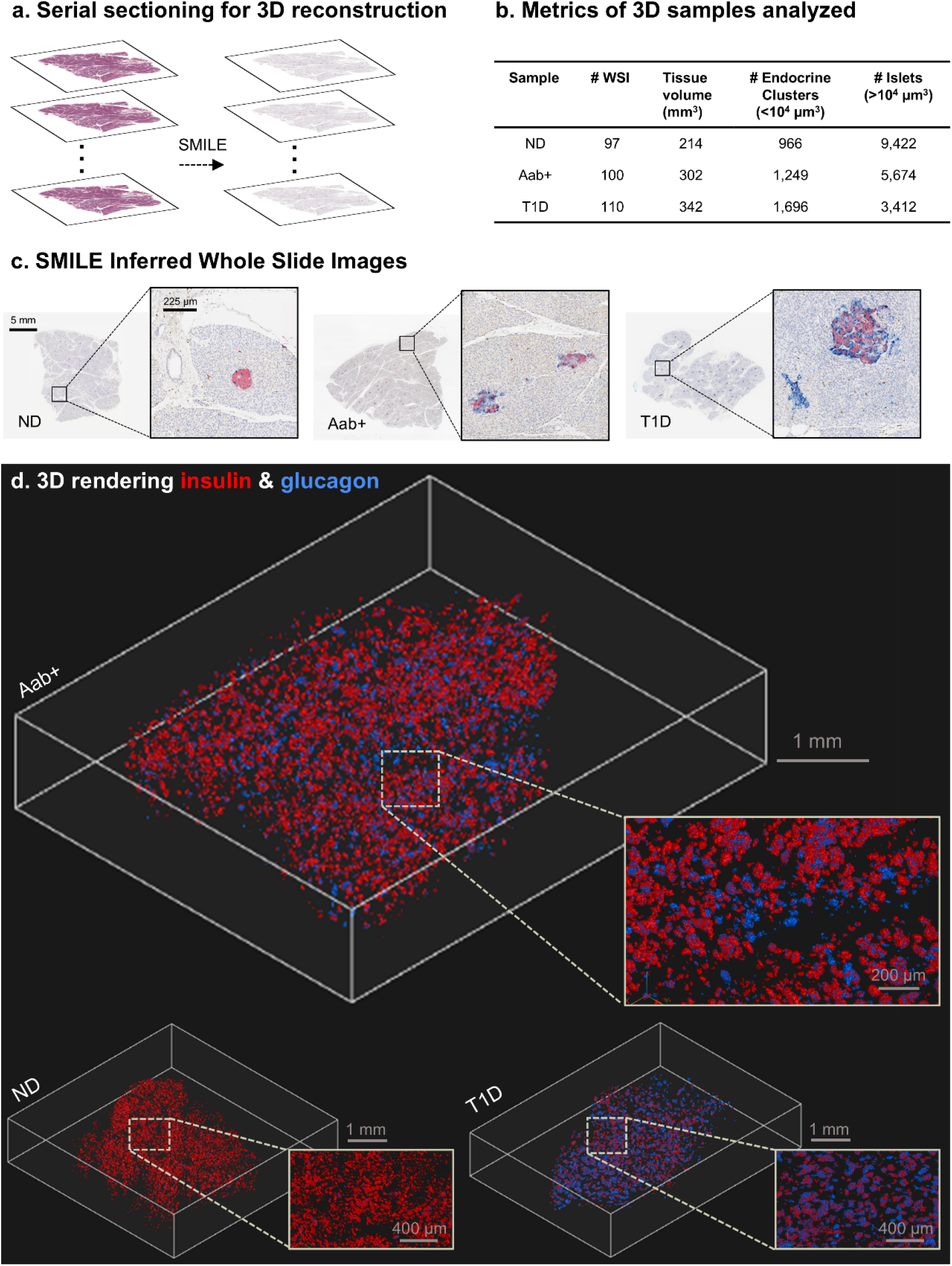
Virtual labeling and 3D reconstruction of normal and diabetic pancreatic islets. **(a)** Serial H&E images were converted to mIHC for 3D islet reconstruction using SMILE. **(b)** Table depicting metrics of the three 3D reconstructed volumes, collected from one nondiabetic (ND) donor (ND), one donor positive for auto-antibodies (Aab+), and one type 1 diabetic (T1D) donor. (c) Sample whole slide mIHC images inferred from serial H&E from each 3D dataset. **(d)** 3D renderings of the inferred insulin and glucagon in large 3D samples of human pancreas tissue depict the dramatic loss in insulin content with type 1 diabetes onset.

Visualization of the resulting inferred WSIs revealed convincing mIHC pathology (**Fig 4c**), with large regions of negative antibody label (but visible, non-endocrine pancreatic parenchyma) interspersed with sparse, round islets labeled positively with insulin (in red) and glucagon (in blue). The islets in the ND sample appeared visibly richer in insulin, with less insulin and more CD3 (in brown) in the Aab+ and T1D histology. 3D renderings confirmed these 2D observations, with visibly redder, insulin-positive islets in the ND donor compared to the Aab+ and T1D (**Fig 4d**). In 3D, the islets appeared as small, variably sized, semi-round objects, consistent with expectations based on 2D observations and previous 3D imaging studies.^49^

We further performed numerical evaluation of the generated 3D volumes (**Fig 5a**). First, we compared the density and size of endocrine clusters and islets in the samples. We found that the density of endocrine clusters between ND, Aab+, and T1D was roughly similar, but that islet density decreased dramatically in Aab+ and T1D sample volumes. Additionally, we found significantly increased mean islet size with T1D progression, and increasing variability in islet size, suggesting that pancreatic tissue heterogeneity increases with T1D onset. Next, we compared the composition of the insulin and glucagon within, and the CD3 signal in the 50 µm around each endocrine cluster (**Fig 5b**) and islet (**Fig 5c**). We found that, as expected, relative insulin composition in the endocrine clusters and islets decreased with progression from ND to AaB+ to T1D, while the CD3 signal increased, reflecting peri-islet inflammation. Further, we found that even with full T1D onset, some islets with high insulin content remained, highlighting the spatial heterogeneity of insulin loss with T1D progression. These findings build on complementary findings in previous works in T1D,^49,58,59^ validating the biological relevance of our virtual labeling approach.

**Figure 5:**
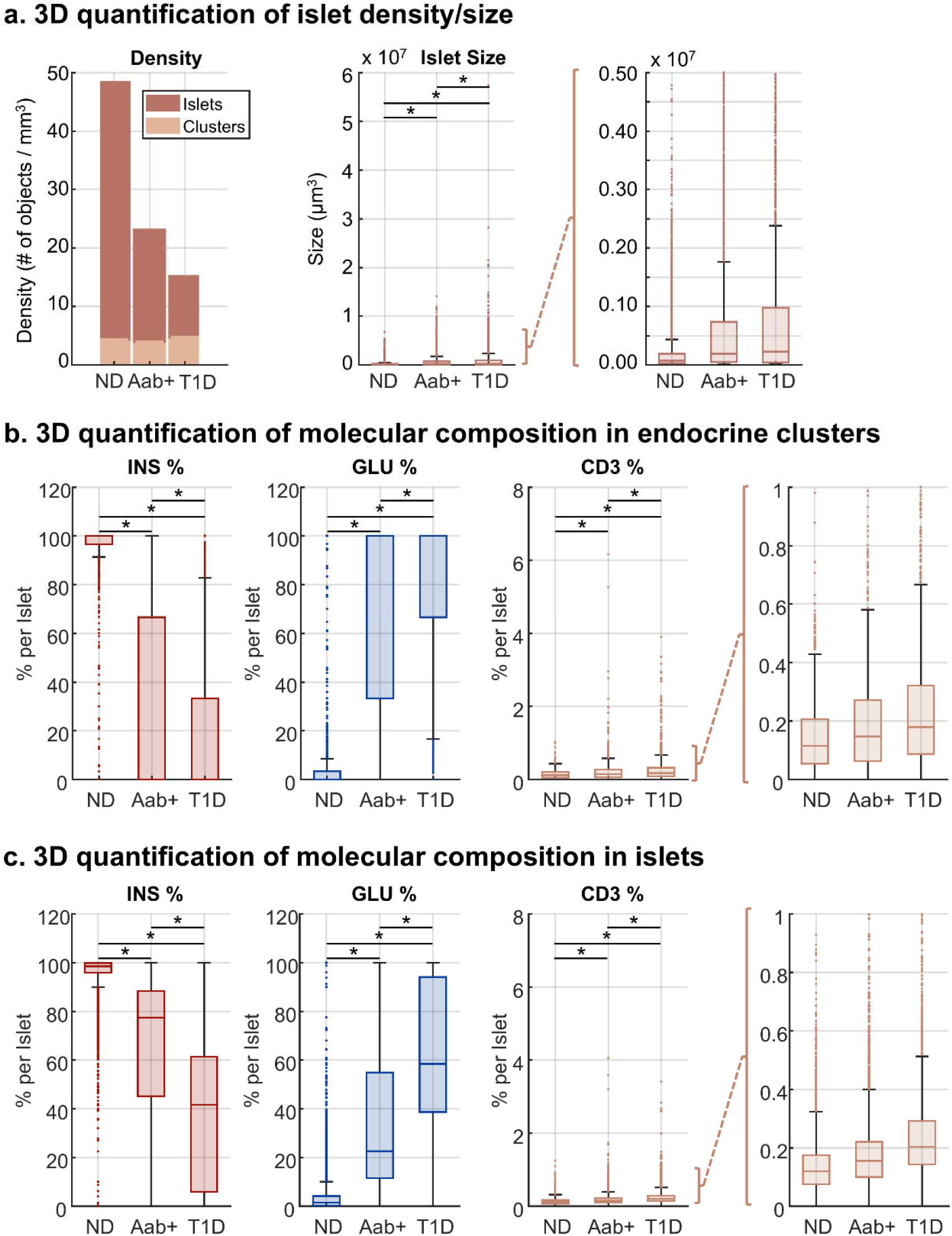
3D Quantitative, spatial analysis of pancreatic islets. **(a)** left: bar graph depicting endocrine cluster (volume <10^4^ µm ^3^) and islet (volume >10^4^ µm^3^) density across ND, Aab+, and T1D tissue. Cluster density is relatively consistent across the samples while islet density decreases from ND to AaB+ to T1D. Middle: boxplot displaying islet size across three samples. The size average islet size and islet size variability significantly increases with T1D progression. Right: zoom in view of the middle plot. **(b)** per-endocrine cluster percent insulin and percent glucagon across ND, Aab+, T1D tissue, highlighting a more heterogenous molecular composition with T1D progression. Percent CD3+ regions within 50 µm of each endocrine cluster depict significantly increasing insulitis with T1D progression. Right: zoom in view of the third plot. **(c)** The same quantifications as in (b) applied to pancreatic islets. For all graphs, * = p < 10 ^-5^using the one-way ANOVA test.

### SMILE stain conversion is highly generalizable

Finally, we evaluated the generalizability of SMILE in two ways. First, we tested the ability of SMILE to infer the triple IHC panel on pancreatic histology collected from surgical resections performed at an independent institution. Second, we applied SMILE to two additional H&E-IHC datasets to assess its performance beyond the pancreatic endocrine system.

First, we evaluated the generalizability of the SMILE workflow using an external dataset. For this test, we retrospectively collected tissue from four patients who underwent surgical resection for pancreatic ductal adenocarcinoma (PDAC), the most common type of pancreatic cancer. Two of the patients were confirmed nondiabetic, while two were previously diagnosed with T1D. With this cohort we tested SMILE’s ability to generalize to H&E images collected at an independent site, as well as SMILE’s ability to infer mIHC on pancreatic tissue affected by nearby cancer, as our training dataset consisted entirely of organ donor tissue. Overall, this tissue appeared more fibrotic and possessed a different H&E appearance as it was processed, stained, and scanned using different protocols from our training cohort. For each of the four tissue specimens, we generated two serial sections. The first section was stained with H&E and digitized, and SMILE was applied to infer virtual mIHC images (**Fig S6**). The second serial section was stained with an mIHC panel for insulin-glucagon-CD3 following the same protocol performed at the original site. While the independent mIHC images contained some non-specific staining, they were of sufficient quality for visual comparison to our virtual mIHC results. The results of SMILE show similar high-quality inference of islet morphology to our internal testing cohort, with high insulin content in the ND cases, and lower insulin, higher glucagon, and higher CD3 content in the T1D cases. Additionally, we find correct inference of antibody negative regions of the images including the pancreatic exocrine compartment and regions of the images previously unseen by the model in the organ donor tissue used for training, such as locations containing fibrosis or mucinous pancreatic cysts. Specifically, islet regions identified in the virtual mIHC output corresponded well to the ground truth locations, and marker co-localization showed high concordance with available reference data. These findings confirm the SMIL model’s capability to generalize to data from external institutions and suggest strong potential for its broader applicability in diverse clinical and research settings.

Second, we tested the SMILE workflow adaptability on single-marker IHC using a public dataset.^60^ We trained and tested SMILE, p2p, and p-p2p models on paired H&E and adjacent IHC sections stained for HER2 and Ki67. Visual inspection of the virtual outputs revealed that all three models successfully learned to generate cellular and structural IHC patterns from the H&E inputs, though virtual image quality varied across models and markers (**Fig 6a**). SMILE demonstrated superior performance in recapitulating both membranous HER2 staining pattern and nuclear Ki67 expression, preserving staining intensity and spatial distribution more accurately than p2p and p-p2p approaches. Quantitative evaluation across seven metrices (SSIM, CW-SSIM, PSNR, FID, CMMD, VMMD, UNI2) confirmed SMILE’s consistent superiority, with radar plots showing larger coverage areas for both HER2 and Ki67 stains. The p2p model showed intermediate performance, outperforming SMILE only on metrics such as SSIM and PSNR which compare pixel-to-pixel similarity and do not accurately sample the spatial morphology of the histology, while p-p2p exhibited more variable result with occasional loss of staining specificity and reduced contrast in virtual outputs (**Fig 6b**).

**Figure 6:**
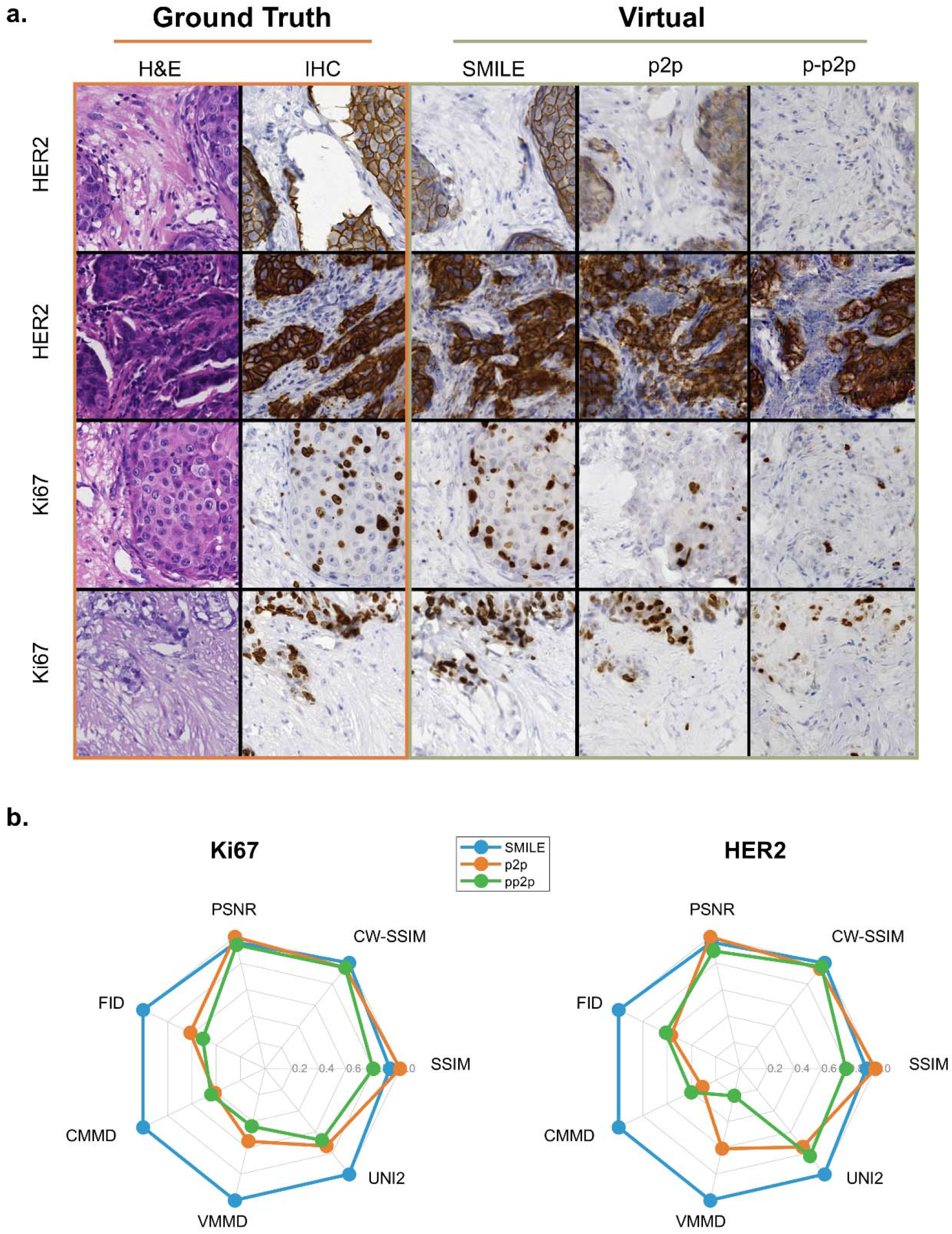
Visual comparison of ground truth and virtual tiles from public dataset. **(a)** Mosaic representing ground truth histology (GT H&E, GT IHC) and virtual IHC generated using three models (SMILE, p2p, p-p2p) of four representative testing tiles from two kinds (HER2, Ki67) of IHC stains. **(b)** radar plots representing quantitative evaluation results using seven methods (SSIM, CW-SSIM, PSNR, FID, CMMD, VMMD, UNI2) for Ki67 (left) and HER (right) stains. Larger polygon areas indicate superior model performance. SMILE (blue) consistently outperforms p2p (orange) and p-p2p (green) across multiple metrices for both markers.

## DISCUSSION

Our virtual labeling workflow addresses the critical time and labor limitations of conventional mIHC. In this work, we developed and validated a diffusion model-based virtual labeling pipeline to transform whole slide H&E images into triple-plex mIHC labeling insulin, glucagon, and CD3. Using a curated dataset of 212 registered WSI pairs from 72 donors spanning diverse diabetes-relevant demographics, we revealed that SMILE significantly outperforms conventional GAN-based architectures in both quantitative matrices and human visual perception. We successfully extended this workflow to whole-slide inference and 3D volumetric reconstruction, which provide both a functional tool and a roadmap for developing virtual labeling technologies in multidimensional molecular analysis from routine H&E slides.

A major contribution of this study work is our strategic dataset construction addressing the severe class imbalance inherent to pancreatic tissue, where islets constitute only 1-5% of total mass.^56,57^ Our islet-centered sampling strategy, coupled with rigorous quality control, ensured balanced representation while maintaining biological diversity and registration fidelity across more than 40,000 tile pairs. The inclusion of tissue from diverse donor demographics further enhances translational potential and may be used as a valuable benchmarking dataset for novel virtual labeling methods.

Our comparative evaluation revealed that SMILE consistently outperformed p2p and p-p2p across the majority of metrices, particularly in distribution-based assessments that capture global perceptual quality and biological realism. However, traditional metrics may not perceptually evaluate the realistic or clinically useful virtual labels.^61,62^ These observations align with our findings. While p2p achieved marginally higher pixel-level similarity scores, SMILE demonstrated substantial advantages in clinically relevant metrices. Matrices that leverage pretrained foundation models can more accurately evaluate semantic and contextual fidelity rather than mere pixel-level correspondence, which is a critical distinction given the inherent spatial misalignment between adjacent tissue sections. The label-based metrices further corroborated SMILE’s biological accuracy, with lower regional label differences and reduced global insulin and glucagon differences indicating faithful recapitulation of endocrine cell marker distributions. Pathologist assessments independently validate these findings, with SMILE generated images rated significantly higher for both histological authenticity and concordance with ground truth mIHC.

SMILE introduces several key technical innovations to virtual labeling workflow. Notably, our single-stage diffusion model employs learned drift functions tailors to bridge the complex H&E-to-tiple-IHC distribution, enabling direct conversion of routine H&E slides into multiplexed molecular labels. Furthermore, optimizing our workflow for fp16 image conversion as well as intaking optimal model weight and NFE value during inference process can efficiently enhance SMILE workflow while maintaining superior performance, facilitating scalable whole-slide and volumetric inference. In addition, the SMILE workflow also demonstrates reliable performance on external samples from a different institution. Together, these advances set a new benchmark for virtual mIHC methodologies, catalyzing broader adoption and further development in multidimensional molecular tissue analysis in multicenter studies.

The application of virtual labeling to serial sections enabled unprecedented 3D visualization of endocrine cell distributions across type 1 diabetes progression from H&E images alone. Reconstructed volumes revealed progressive architectural disruption consistent with established T1D pathophysiology, including increased heterogeneity in islet cellular composition and reduced volumetric uniformity reflecting beta cell loss and immune infiltration. The ability to generate coherent 3D representations from virtually labeled serial H&E section with matching patterns observed in actual mIHC confirms the biological accuracy and spatial consistency of SMILE. This validation through 3D reconstruction establishes confidence in the method’s performance for both 2D tissue analysis and more complex volumetric studies. Virtual mIHC labeling addresses critical limitations of conventional mIHC by enabling molecular profiling from H&E sections. However, several limitations of this workflow need to be considered. Our training dataset consists of registered, pseudo-paired adjacent tiles rather than histological labels performed on the same tissue section, which introduces uncertainty in pixel-level correspondence or single cell level accuracy. Second, while powerful, SMILE is computationally intensive, with inference of a single WSI taking approximately 1 hour on a deep learning-capable computer. Future work to improve the inference speed of this or similar Schrödinger bridge models is imperative to improve their broad utility to the field.

## CONFLICT OF INTEREST STATEMENT

The authors declare no conflicts of interest.

## Acknowledgements

The authors acknowledge the following sources of support: JHU Oncology Tissue & Imaging Services (OTIS); U54CA268083; Rolfe Pancreatic Cancer Foundation; Lustgarten Foundation-AACR Career development award for pancreatic cancer research in honor of Ruth Bader Ginsburg; Break Through Cancer Data Science; The Stringer Foundation; The Joseph C. Monastra Foundation for Pancreatic Cancer Research; The Carol and Robert Long Fund for Pancreatic Cancer Research; The Carl Nale Fund for Pancreatic Cancer Research; The Sol Goldman Pancreatic Cancer Research Center; Susan Wojcicki and Denis Troper; Fight Cancer Stay Positive.

## Code Availability

The original implementation of the I^2^SB algorithm is available at the following github: https://github.com/NVlabs/I2SB. The SMILE algorithm with adaptations to the I^2^SB implementation to improve inference time and enable whole slide inference will be made available following peer-reviewed publication of this manuscript.

## Data Availability

The registered, whole slide images and H&E-mIHC tile pairs will be made available following peer-reviewed publication of this manuscript.

## SUPPLEMENTARY TABLES AND FIGURES

**Supplementary Table 1:** Summary of paired and weakly paired virtual labeling/image-to-image translation methods in histopathology. Base models are categorized as GANs, Diffusion, or Schrödinger Bridge (SB). The “From” and “To” columns indicate the source and target image domains for image-to-image translation. The heterogeneity of quantitative evaluation metrics across studies highlights the lack of a standardized, human-perception-oriented metric in the field.

| Author | From | To | Base model | Quantitative evaluation |
| --- | --- | --- | --- | --- |
| Wang et al.<sup>14</sup> | Unlabeled | H&E | GANs | NRMSE, NMI, PSNR, MSSIM |
| Li et al.<sup>15</sup> | Unlabeled | H&E, PSR, Orcein | GANs | None |
| Zhang et al.<sup>16</sup> | Unlabeled | H&E, PSR, EVG | GANs | SSIM, PSNR, LPIPS |
| Rana et al.<sup>17</sup> | Unlabeled | H&E | GANs | PCC, PSNR, SSIM |
| Khan et al.<sup>18</sup> | Unlabeled | H&E | GANs | PCC, PSNR, SSIM |
| Liu et al.<sup>26</sup> | H&E | IHC | GANs | PSNR, SSIM |
| Rivenson et al.<sup>19</sup> | Unlabeled | H&E, JS, MT | GANs | None |
| Ghahremani et al.<sup>25</sup> | IHC | mpIF | GANs | AJI, Dice |
| Li et al.<sup>20</sup> | Unlabeled | H&E | GANs | SSIM, PSNR |
| Ounissi et al.<sup>21</sup> | H&E | IHC | GANs | MSE, PSNR, SSIM |
| Li et al.<sup>22</sup> | H&E | IHC | GANs | SSIM, PHV, FID, KID |
| Peng et al.<sup>23</sup> | H&E | IHC | GANs | SSIM, FID, DISTS |
| Chen et al.<sup>24</sup> | H&E | IHC | GANs | mIOD, Pearson-R, PSNR, SSIM |
| Kataria et al.<sup>28</sup> | H&E | IHC | Diffusion | SSIM, PSNR, FID, KID |
| He et al.<sup>29</sup> | H&E | IHC | Diffusion | SSIM, PSNR, FID, VIF, Hist |
| Zhang et al.<sup>31</sup> | Unlabeled | H&E | Diffusion | SSIM, PSNR, LPIPS |
| Abraham & Levenson<sup>33</sup> | SFM | H&E | Diffusion | FID |
| Zheng et al.<sup>32</sup> | Polarimetric | H&E, Fluor | Diffusion | PSNR, SSIM, FID, LPIPS |
| Yang et al.<sup>30</sup> | H&E | IHC | Diffusion | DISTS, FID, KID, PSNR, SSIM |
| Qiu et al.<sup>36</sup> | H&E | IHC | SB | PSNR, SSIM, FID, KID |

**Supplementary Table 2.** Cohort Demographics. describing the cohort size including aggregate sex, age, race, anatomical location of resection within the pancreas, and diabetes status.

| **Description** | **# donors** |
| --- | --- |
| **Cohort** |  |
| Initial cohort (donor) | 72 |
| External cohort (surgery) | 4 |
| **Sex** |  |
| Male | 45 |
| Female | 31 |
| **Age (years)** |  |
| < 10 | 15 |
| 11 – 20 | 25 |
| 21 – 30 | 30 |
| 31 – 40 | 2 |
| 41 – 50 | 0 |
| 51 – 60 | 2 |
| > 60 | 1 |
| **Donor race** |  |
| Caucasian | 49 |
| Black | 9 |
| Hispanic | 14 |
| Other | 1 |
| Unknown | 4 |
| **Location of pancreatic resection** |  |
| Head | 71 |
| Body | 70 |
| Tail | 74 |
| Unknown | 1 |
| **Diabetes status** |  |
| Without diabetes | 26 |
| Auto-antibody positive | 24 |
| Type 1 diabetic | 26 |

**Supplementary Figure 1:**
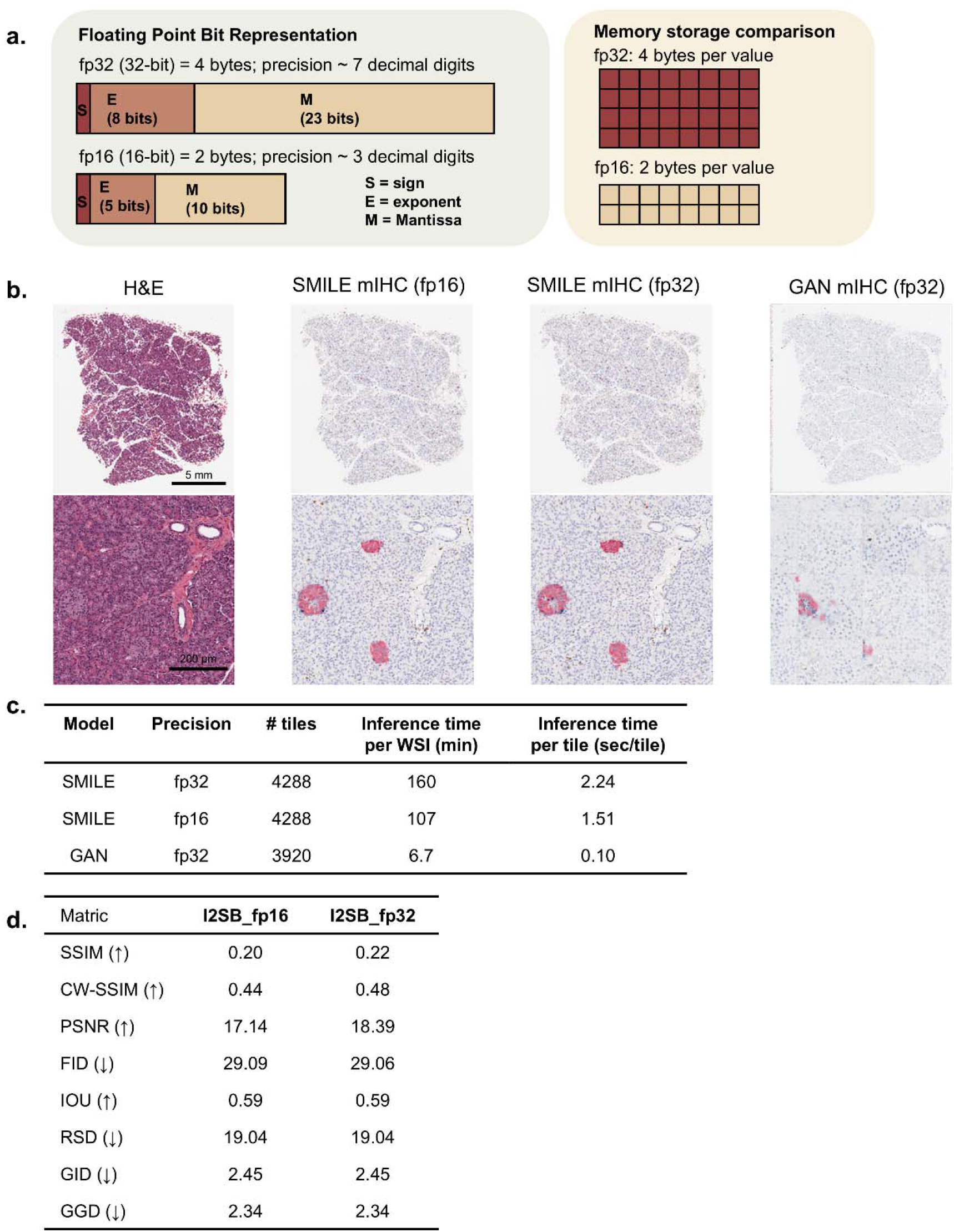
performance evaluation of reduced-precision representation in virtual labeling. **(a)** schematic representation of fp32 and fp16 floating-point bit along with memory compression from 4 bytes to 2 bytes per value. **(b)** representative H&E-stained tissue sections and corresponding virtual mIHC results from SMILE and GAN models at different precision levels (fp16 and fp32). **(c)** Computational performance comparison across models showing inference time per tile. GAN demonstrated superior computational efficiency. Reducing image precision significantly improved SMILE efficiency. **(d)** Image quality matrices comparing three model performances. Matrices indicate minimal quality degradation with fp16 precision while achieving 50% memory reduction.

**Supplementary Figure 2:**
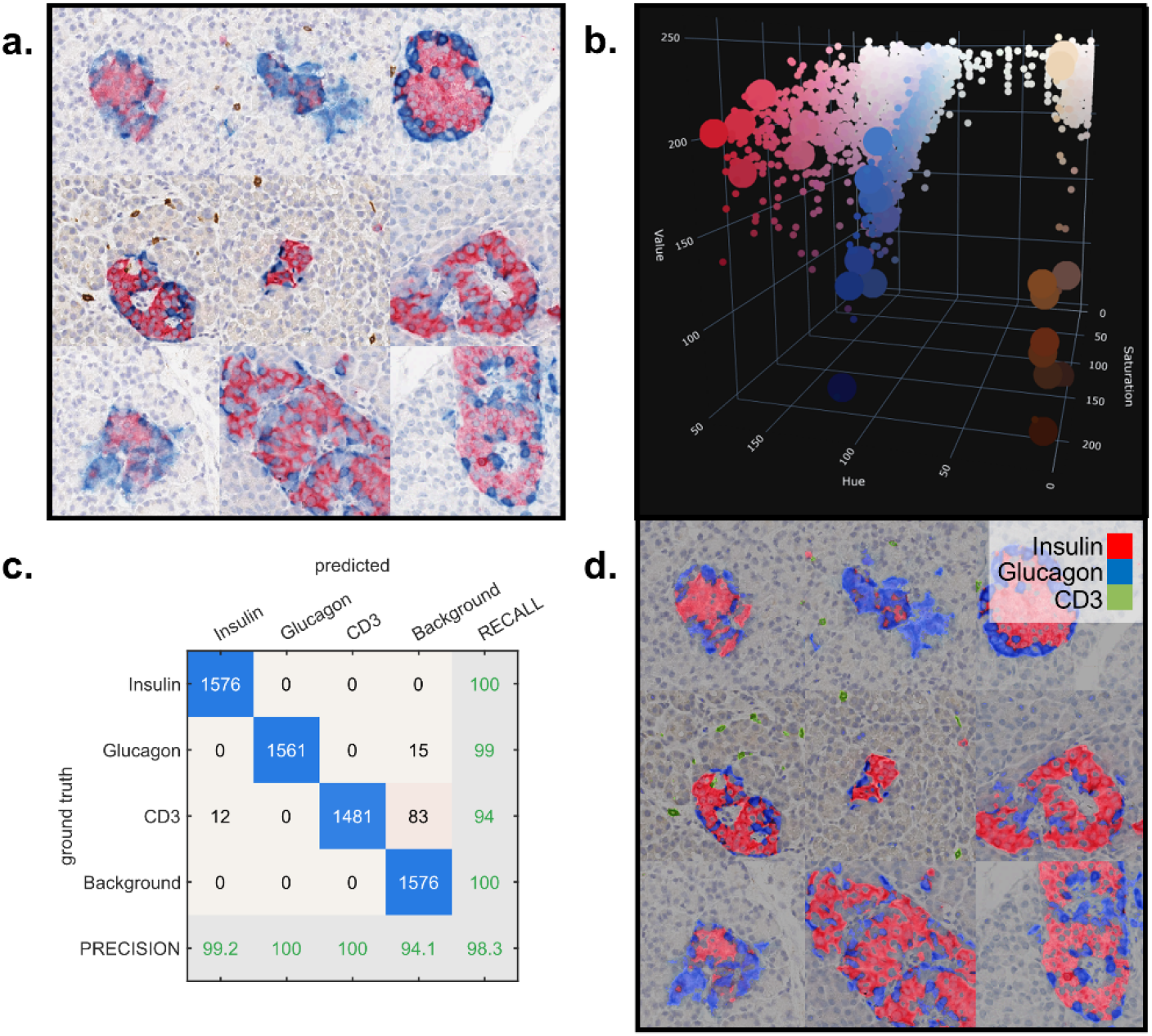
Label mask optimization using hue-saturation-variation (HSV) color space. **(a)** A 3 × 3 mosaic image containing islet from mIHC images from nine independent donors. **(b)** Colorized 3D scatter plot depicting the HSV value of each pixel in the mosaic image shown in (a). Using this visualization, HSV thresholds were determined to delineate the unlabeled, insulin+, glucagon+, and CD3+ pixels. **(c)** Confusion matrix depicted HSV segmentation performance compared against manual annotation of an independent mosaic image. **(d)** HSV-based segmentation mask of the mosaic image shown in (a).

**Supplementary Figure 3:**
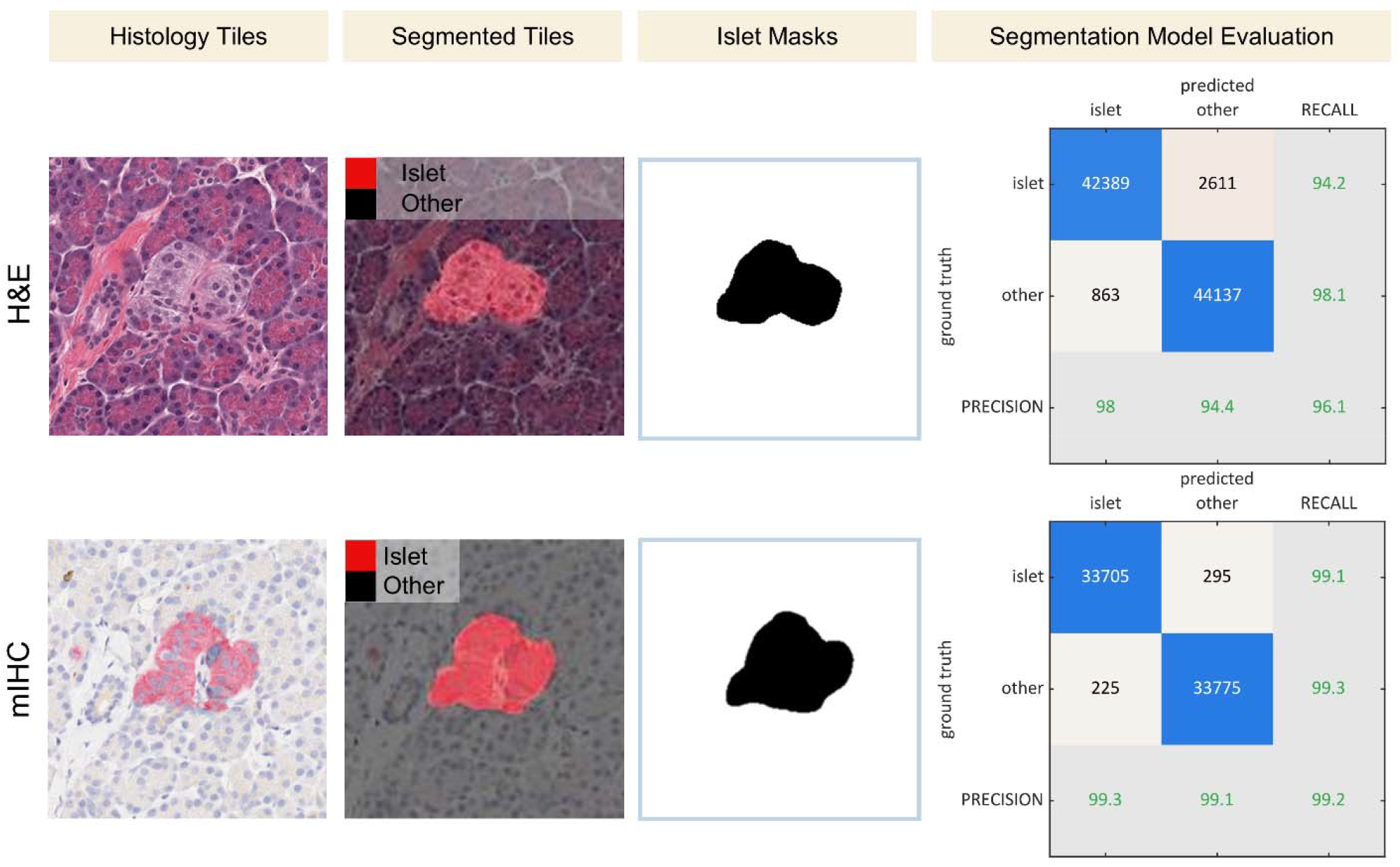
islet mask generation using CODA method. Deep learning based semantic segmentation masks of pancreatic islets were trained in H&E and mIHC to determine islet morphology. Confusion matrices for both models provided.

**Supplementary Figure 4:**
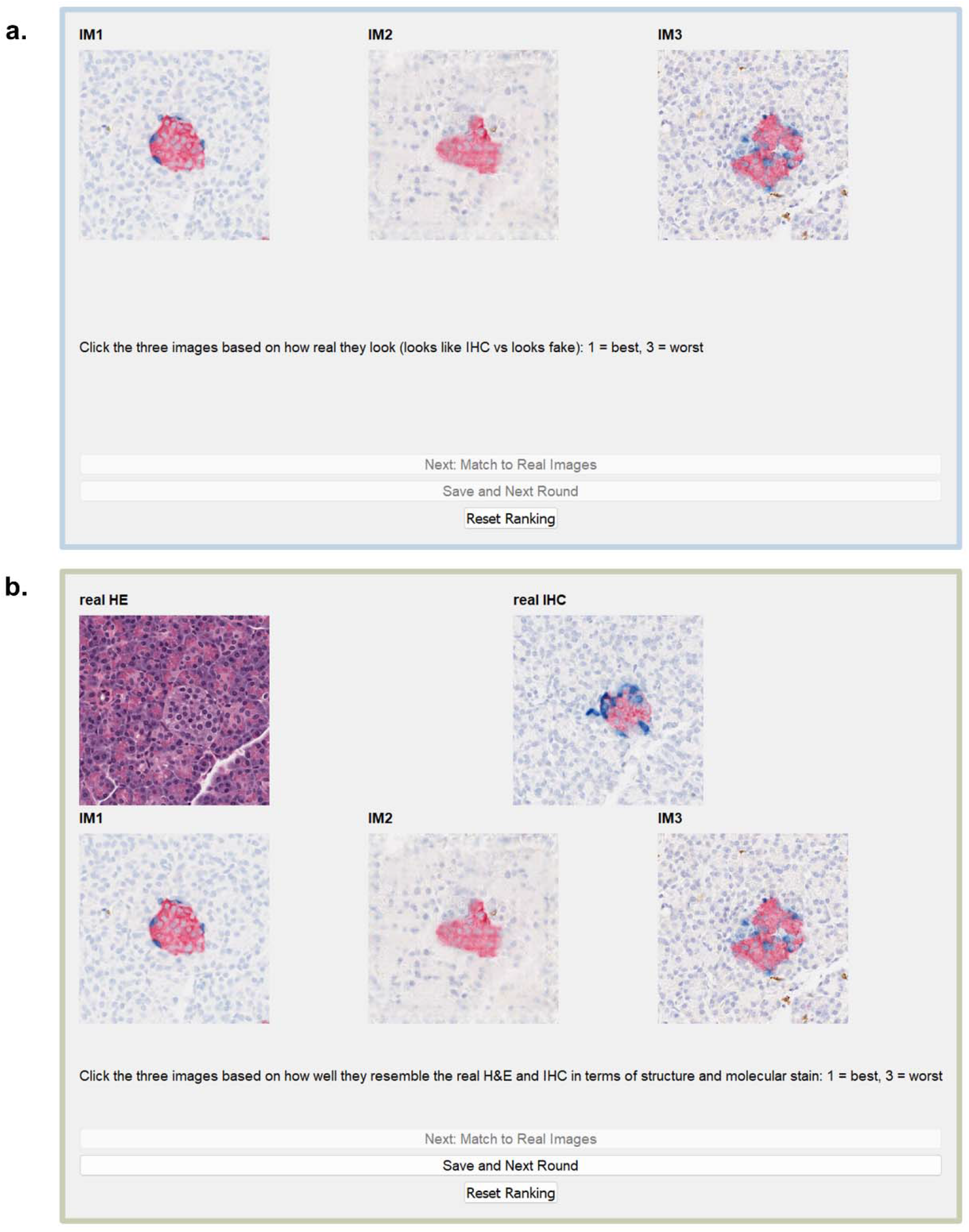
Graphical user interface (GUI) for blinded tile review by pancreatic pathologists. **(a-b)** Sample screenshot taken from a GUI designed to solicit blinded feedback on the virtual mIHC images. **(a)** The first page of the GUI showed the user three virtually generated mIHC images (one made by p2p, one by p-p2p, and one by SMILE) and asked the user to rank based on which image looked the most real. **(b)** The second page of the GUI showed the user the same three virtual images, but now also showed the corresponding ground truth (GT) H&E and GT mIHC. The GUI prompted the user to rank the three virtual images based on their ability to accurately capture the islet morphology of the GT H&E and the molecular labeling of the GT mIHC image.

**Supplementary Figure 5:**
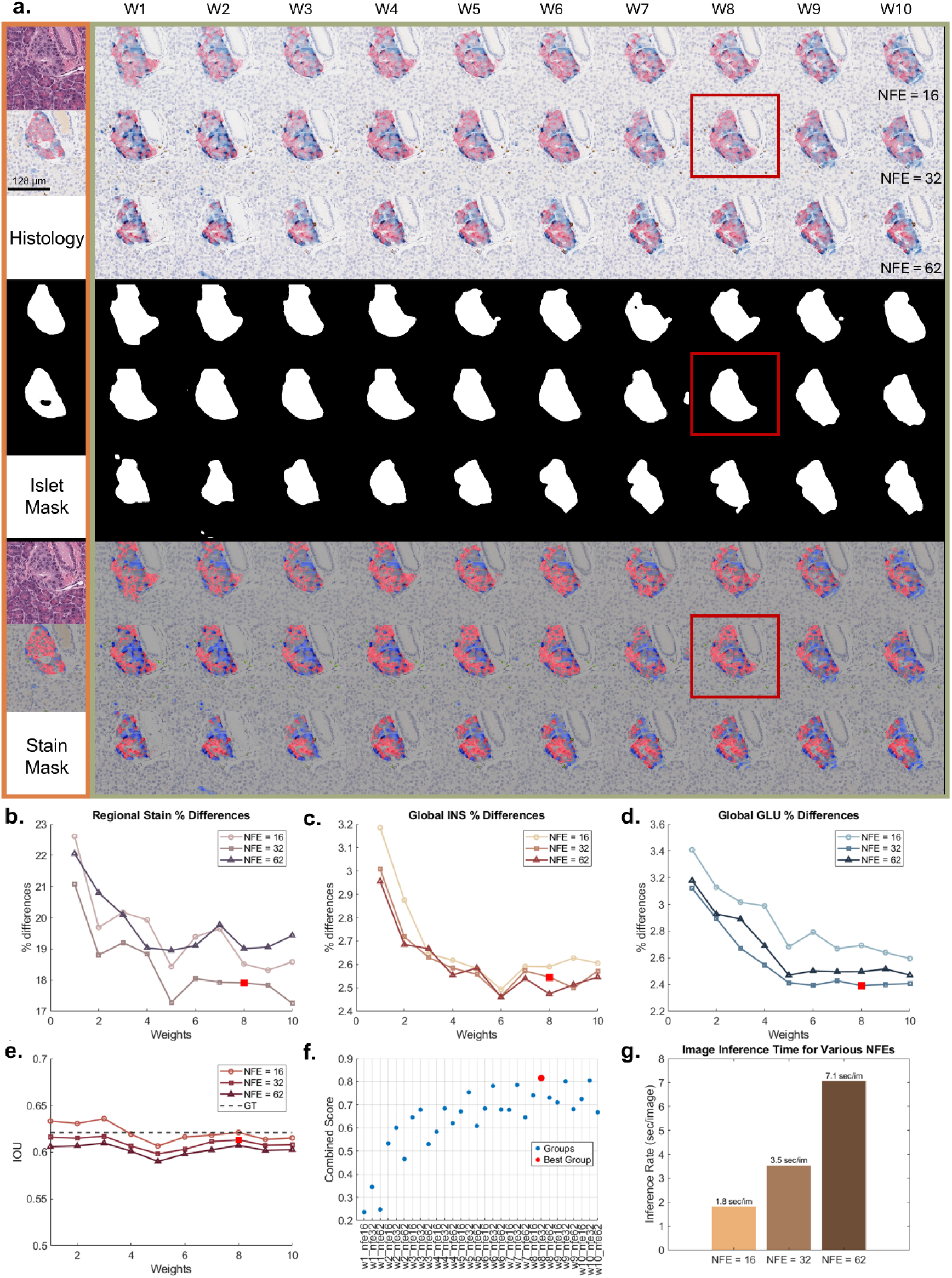
Determination of optimal model weights and number of function evaluations (NFE) for SMILE inference: **(a)** Image mosaic depicting sample histology, islet masks, and label masks of GT H&E and mIHC tiles and with SMILE inference results from ten model weights and three different NFEs. The islet and label masks were used to calculate the label composition and islet morphological differences for each weight and each NFE **(b - e)**. Combining the scores obtained from these calculations revealed the optimal combination (weight = 8, NFE = 32). **(g)** depicts the SMILE inference time for a 256 × 256 tile for the three different NFEs.

**Supplementary Figure 6:**
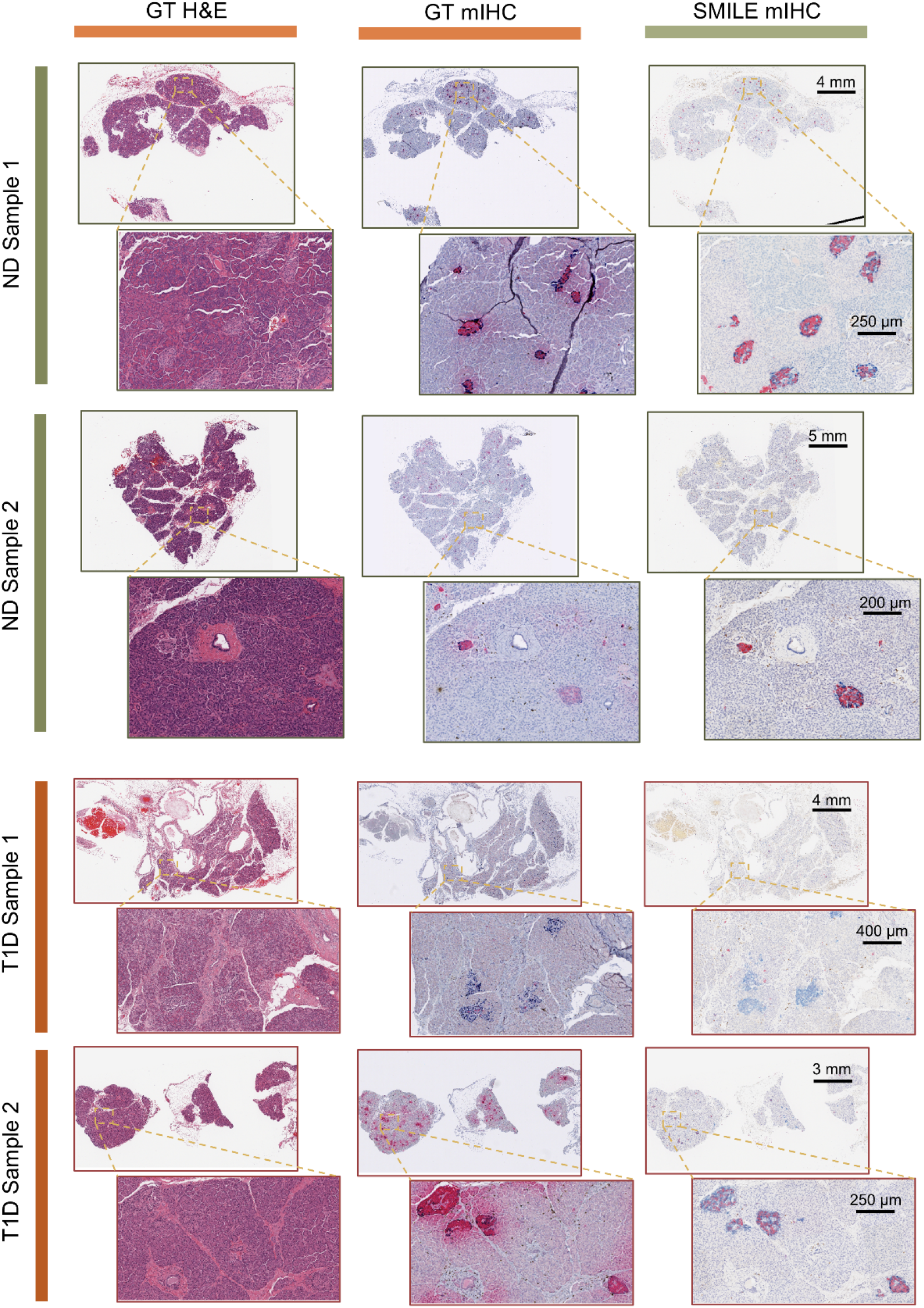
Test generalizability of SMILE workflow by analyzing pancreatic samples from an external institution. Representative images from two T1D (upper two panels) and two ND samples (lower two panels) are shown. The left column displays GT H&E slides at both WSI and zoomed-in views. The middle column displays virtual mIHC slides at WSI and zoomed-in views. The right column displays GT mIHC slides, which are adjacent to the GT H&E slides and were stained by JHU technicians.

## SUPPLEMENTARY MATERIALS

### SUPPLEMENT 1: TYPES OF GENERATIVE MODELS USED HERE

This supplement provides technical details of the generative models used in our study. For training and sampling, we used the default hyperparameters for p2p (https://github.com/matlab-deep-learning/pix2pix), p-p2p (https://github.com/bupt-ai-cz/BCI), and I^2^ SB (https://github.com/NVlabs/I2SB).

### Generative Adversarial Networks

We trained two conditional GAN (cGAN) models: pix2pix (p2p) ^13^ and pyramid-pix2pix (p-p2p).^26^ GANs consist of two networks trained adversarially: a generator *G* that synthesizes images and a discriminator *D* that distinguishes real from generated samples. The standard GAN objective is formulated as a minimax game:

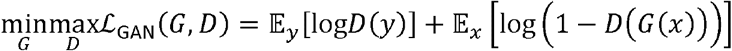

where *x* is the source image and *y* is the target image. For paired image-to-image translation, the conditional GAN (cGAN) modifies this objective by conditioning both *G* and *D* on the input:

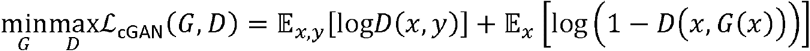

The p2p model additionally incorporates an *L*_1_ reconstruction loss to encourage pixel-wise similarity:

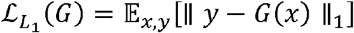

yielding the combined objective 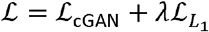. The p-p2p model extends this framework with a multi-scale pyramid architecture for improved performance on high-resolution histopathology images.

While cGANs perform direct image-to-image translation, their adversarial training can lead to instability and mode collapse. Furthermore, the generator architecture fundamentally limits the learned mapping to deterministic transformations, which may not capture the full variability of labeling patterns.

### Denoising Diffusion Probabilistic Models

Denoising diffusion probabilistic models (DDPMs)^27^ define a forward Markov process that gradually corrupts data x_0_ ∼ *p*_0_ by adding Gaussian noise over *T* timesteps:

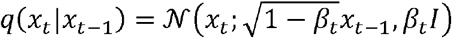

where 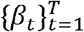 is a variance schedule. The marginal distribution at time *t* can be written as 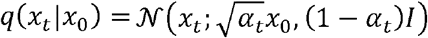 where 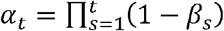. As 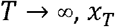 converges to 𝒩 (0, *I*).

Image generation proceeds by learning the reverse process *p*_*θ*_ (*x*_t − 1_|*x*_*t*_) that denoises samples from the prior *p*_*T*_ = 𝒩 (0, *l*) back to the data distribution. The model is trained to predict the noise *ϵ* added at each step, minimizing:

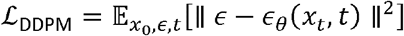

For conditional generation, the denoiser can be conditioned on auxiliary information *y* (e.g., the source image), yielding *ϵ*_*θ*_ (*x*_*t*_, *t, y*). However, conditional DDPMs still initialize sampling from Gaussian noise *x*_*T*_ ∼ 𝒩 (0, *l*), using the conditioning signal only to guide the reverse trajectory. This is fundamentally different from direct image-to-image translation: the source image information is incorporated as a conditioning input rather than serving as the starting point of the generative process. Consequently, conditional DDPMs may not preserve fine structural details from the source domain, limiting their effectiveness for virtual labeling where anatomical correspondence is critical.

### Schrödinger Bridges

The Schrödinger bridge problem, originally formulated by Schrödinger in 1931,^63^ provides a principled framework for finding stochastic processes that transport one probability distribution to another. Unlike DDPMs which map noise to data, Schrödinger bridge methods solve for the optimal path measure connecting two arbitrary endpoint distributions *p*_0_ (source) and *p*_1_ (target).

The Schrödinger bridge problem seeks the path measure ℚ^*^ closest to a reference process ℙ (typically Brownian motion) while satisfying the boundary constraints:

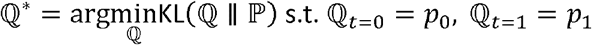

This formulation is equivalent to entropy-regularized optimal transport, providing a geometrically meaningful interpolation between distributions.

The Image-to-Image Schrödinger Bridge (I^2^ SB^36^) implements this framework for paired image translation.

Given paired training data (*x*_0_, *x*_1_) ∼ π (*x*_0_, *x*_1_) where *x*_0_ is the source (H&E) and *x*_1_ is the target (IHC), I ^2^SB learns forward and backward diffusion processes that optimally transport between the marginals. The forward process interpolates between the paired samples:

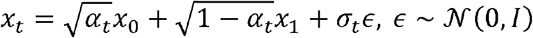

where *σ*_*t*_ controls the stochasticity. A neural network is trained to predict the target *x*_1_ from intermediate states, enabling sampling that starts directly from source images *x*_0_ rather than Gaussian noise.

This formulation offers key advantages for virtual labeling: (1) the source image serves as the initialization, preserving structural information throughout the translation; (2) the optimal transport framework ensures geometrically consistent mappings that minimize unnecessary distortion; and (3) the model learns the joint coupling between source and target distributions rather than conditional generation from a fixed prior. These properties make Schrödinger Bridge methods theoretically well-suited for paired image translation tasks where maintaining correspondence between input and output is essential.

During inference, the number of function evaluations (NFE) can be reduced with minimal quality degradation due to the optimal transport structure, enabling efficient whole-slide image translation.

### SUPPLEMENT 2: IDENTIFY OPTIMAL MODEL WEIGHT AND NUMBER OF FUNCTIONAL EVALUATIONS

We holistically reviewed the insulin, glucagon, and CD3 labels using both the islet masks and the label masks to mathematically compute one beneficial variable, intersection over union, and three detrimental variables, regional insulin difference, global insulin difference, and global glucagon difference. We normalized these variables using min-max normalization and compute the overall score for each model weight and number of functional evaluations combination.

Intersection over union (IOU) formula:

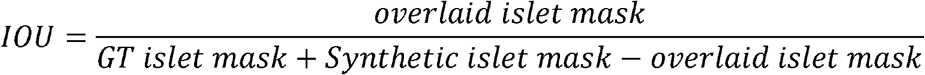

Regional insulin difference (RID) formula:

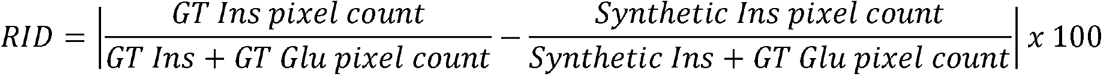

Global insulin difference (GID) formula:

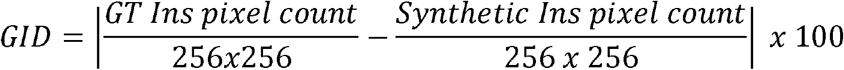

Global glucagon difference (GGD) formula:

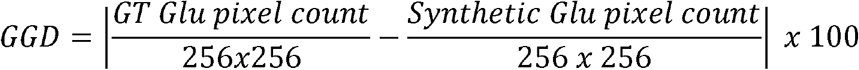

Min-max normalization formula for beneficial variable:

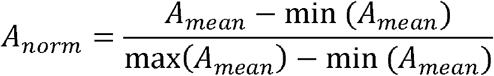

Min-max normalization formula for detrimental variable:

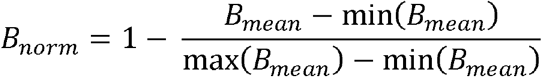

Overall score formula for N variables with z_k_ represents each normalized variable:

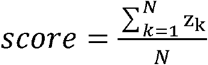

## Notes

### Competing Interest Statement

The authors have declared no competing interest.

